# The Platinum Pedigree: A long-read benchmark for genetic variants

**DOI:** 10.1101/2024.10.02.616333

**Authors:** Zev Kronenberg, Cillian Nolan, David Porubsky, Tom Mokveld, William J. Rowell, Sangjin Lee, Egor Dolzhenko, Pi-Chuan Chang, James M. Holt, Christopher T. Saunders, Nathan D. Olson, Sean McGee, Andrea Guarracino, Nidhi Koundinya, William T. Harvey, W. Scott Watkins, Katherine M. Munson, Kendra Hoekzema, Khi Pin Chua, Cairbre Fanslow, Christine Lambert, Harriet Dashnow, Erik Garrison, Josh D. Smith, Peter M. Lansdorp, Justin M. Zook, Andrew Carroll, Lynn B. Jorde, Deborah W. Neklason, Aaron R. Quinlan, Evan E. Eichler, Michael A. Eberle

## Abstract

Recent advances in genome sequencing have improved variant calling in complex regions of the human genome. However, it is difficult to quantify variant calling performance since existing standards often focus on specificity, neglecting completeness in difficult to analyze regions. To create a more comprehensive truth set, we used Mendelian inheritance in a large pedigree (CEPH-1463) to filter variants across Illumina, PacBio high-fidelity (HiFi), and Oxford Nanopore Technologies platforms. This generated a variant map with over 4.7 million single-nucleotide variants, 767,795 indels, 537,486 tandem repeats, and 24,315 structural variants, covering 2.77 Gb of the GRCh38 genome. This work adds ∼200 Mb of high-confidence regions, including 8% more small variants, and introduces the first tandem repeat and structural variant truth sets for NA12878. As an example of the value of this improved benchmark, we retrained DeepVariant using this data to reduce genotyping errors by ∼34%.

## Introduction

Whole-genome sequencing (WGS) is becoming increasingly common in clinical settings with many laboratories already using it to identify the genetic causes of individuals suffering from rare and undiagnosed genetic diseases (Gonzaga-Jauregui et al., 2012; Marshall, Bick, et al., 2020; Roach et al., 2010; Talkowski et al., 2012). In the future, WGS may also become a common part of an individual’s medical records with specific use cases such as newborn carrier screening (Chen et al., 2020; Punj et al., 2018) and pharmacogenomics testing (Relling & Evans, 2015; van der Lee et al., 2020) likely leading the way. The sequencing pipelines that will be employed may vary significantly between different labs that may utilize different technologies and or bioinformatic workflows. An essential part of implementing WGS in clinical settings is benchmarking the variant calling performance achieved by individual laboratories (Marshall, Chowdhury, et al., 2020). This requires a highly curated set of known “truth” variant calls that labs can easily obtain and process internally to quantify the variant calling performance of their sequencing pipeline.

WGS and variant calling results in millions of variant calls, which require a general way to assess the performance of variant types as a class. Currently, many labs benchmark small variant performance using the Genome in a Bottle (GIAB) standards developed by the National Institute for Standards and Technologies (Wagner et al., 2022; Zook et al., 2016). This standard has been developed using various technologies on either a single sample or trios and represents a highly accurate call set by excluding ambiguous genomic regions. An alternative benchmark, the Platinum Genomes (PG), used Mendelian inheritance in a large, 17-member pedigree to build a highly accurate and more comprehensive small variant truth set (Eberle et al., 2017). Compared to the GIAB resource, the PG benchmark prioritizes sensitivity and completeness at the expense of some specificity. Both of these resources, for NA12878 (second generation female in CEPH-1463), are limited to small variants (single-nucleotide variants [SNVs] and indels) and do not cover more difficult variant types or many complex genomic regions. Recent work has used long-read-based diploid *de novo* assemblies to generate a benchmark for more complex variants, such as tandem repeats (TRs) in the GIAB HG002 sample (English et al., 2024). Compared to a pedigree-based analysis, this benchmark used concordance of assemblies and manual curation to improve the accuracy of the calls, so it excludes indels smaller than 5 bp and the most challenging regions of the genome. As technologies and reference genomes have improved, it is important that standards evolve alongside these technologies to include more of the genome and increasingly complex variant classes (Majidian et al., 2023).

In this study, we sequence an extended family (two parents and eight children) using the latest technologies from Illumina, PacBio, and Oxford Nanopore Technologies (ONT) to create a reference catalog across the spectrum of genetic variation. Identifying the genome-wide inheritance vectors of the parental chromosomes into eight children allows us to adjudicate the pedigree-level genotyping accuracy for SNVs, indels, structural variants (SVs), and TRs. Combining both the pedigree and long reads allows us to deliver a more comprehensive reference dataset that covers both the easy and difficult to analyze parts of the genome. This pedigree-based truth set includes 4,719,711 SNVs, 767,795 indels, 537,486 TRs, and 24,315 SVs that, combined, impact 37,885,857 bp of sequence relative to the GRCh38 reference. In addition to acting as a benchmarking dataset for assessing sequencing pipelines, the data for this study are being made publicly available so that this resource can be used to develop and improve new bioinformatic methods. We demonstrate utility of this dataset by showing that retraining DeepVariant (Poplin et al., 2018) with this data reduces errors in small variant calls by 38.4% for SNVs and 19.3% for indels when evaluated on the Platinum Pedigree truth set, and by 18.2% for SNVs and 5.3% for indels when evaluated on GIAB.

## Results

In a recent study, the four generation CEPH-1463 pedigree was sequenced using multiple technologies and analyzed to study the *de novo* mutation rate for SNVs, indels, TRs, and SVs (Porubsky et al., 2024). For this study, we focus on the haplotypes and variants in G2 that are inherited among the eight children in G3 (**Fig. 1**). While there are ten total samples in G2 and G3, there are only two founders (NA12877 and NA12878) so the genomes of every individual in G3 are a mosaic of the DNA from their parents. The inheritance of the chromosomal segments from the parents provides a framework for identifying the accuracy of variants within this nuclear family at nearly any position within the genome where the inheritance pattern can be identified (Eberle et al., 2017). Importantly, with the exception of ONT ultra-long (UL) data generated by sequencing cell lines, the sequencing data was exclusively derived from blood, thus eliminating the confounding effects of cell line artifacts.

**Fig. 1.**
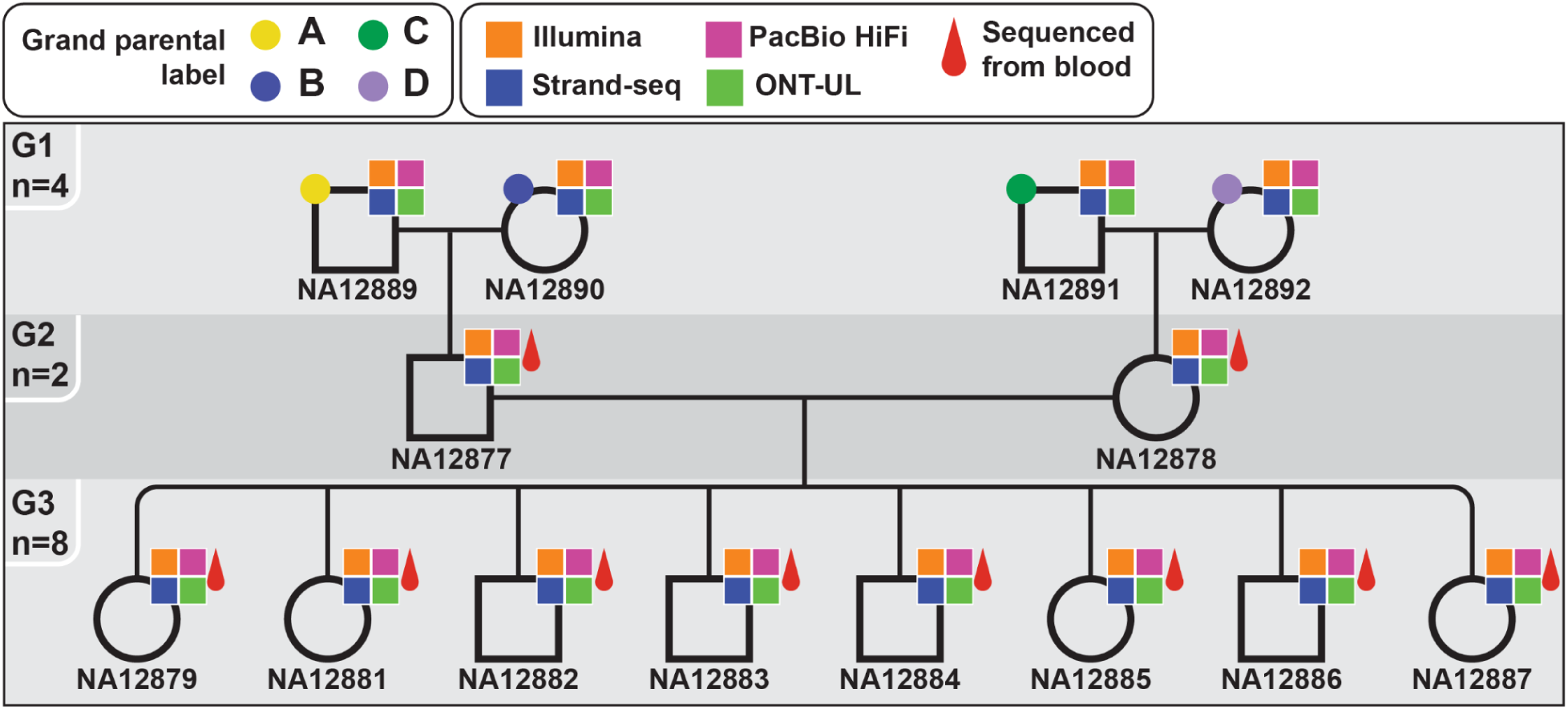
Three-generation CEPH-1463 pedigree and the sequencing technologies applied to each individual. Living individuals were sequenced from blood; the first generation was sequenced from the cell lines. A, B, C & D correspond to the grandparent of origin (A=NA12889; B=NA12890; C=NA12981; D=NA12892). HiFi, high-fidelity; ONT-UL, Oxford Nanopore Technologies ultra-long

### Identification of inheritance vectors

We identified blocks of chromosomal segments transmitted from the parents to each of the eight children using the SNV calls aligned to GRCh38 (**Methods**). For this work, we traced the haplotypes back to the G1 generation and label the chromosomes as A, B, C & D corresponding to the grandparent of origin (A=NA12889; B=NA12890; C=NA12981; D=NA12892). This analysis identified 539 recombination events (**Fig. 2A, Table S1, Fig. S1**) among the eight G3 individuals; 227 (42%) recombination events occurred on paternal (A, B) haplotypes, and 312 (58%) recombination events occurred on maternal (C, D) haplotypes in the autosome. We independently validated 98.1% of the recombination events with Strand-seq data (**Table S2**). Additionally, nine maternal recombinations were identified on Chromosome X. We calculated a 1.29 cM per Mb maternal recombination rate and 0.99 cM per Mb paternal recombination rate, matching previous work in this pedigree and general estimates across human pedigrees (Eberle et al., 2017; Kong et al., 2002). Recombination breakpoints could be refined to a median resolution of 7.1 kb (**Fig. S2**) and we were able to assign 95.2% of the genome to the haplotype of origin. Of the 549 autosomal segments where we have determined the transmission, we observed all four haplotypes (A, B, C & D) in 539 of the segments covering 2.79 Gb of the genome (**Fig. 2B**). The remaining ten regions span 41 Mb, where only three of the four haplotypes were transmitted to G3 (**Fig. S3; Table S3**).

**Fig. 2.**
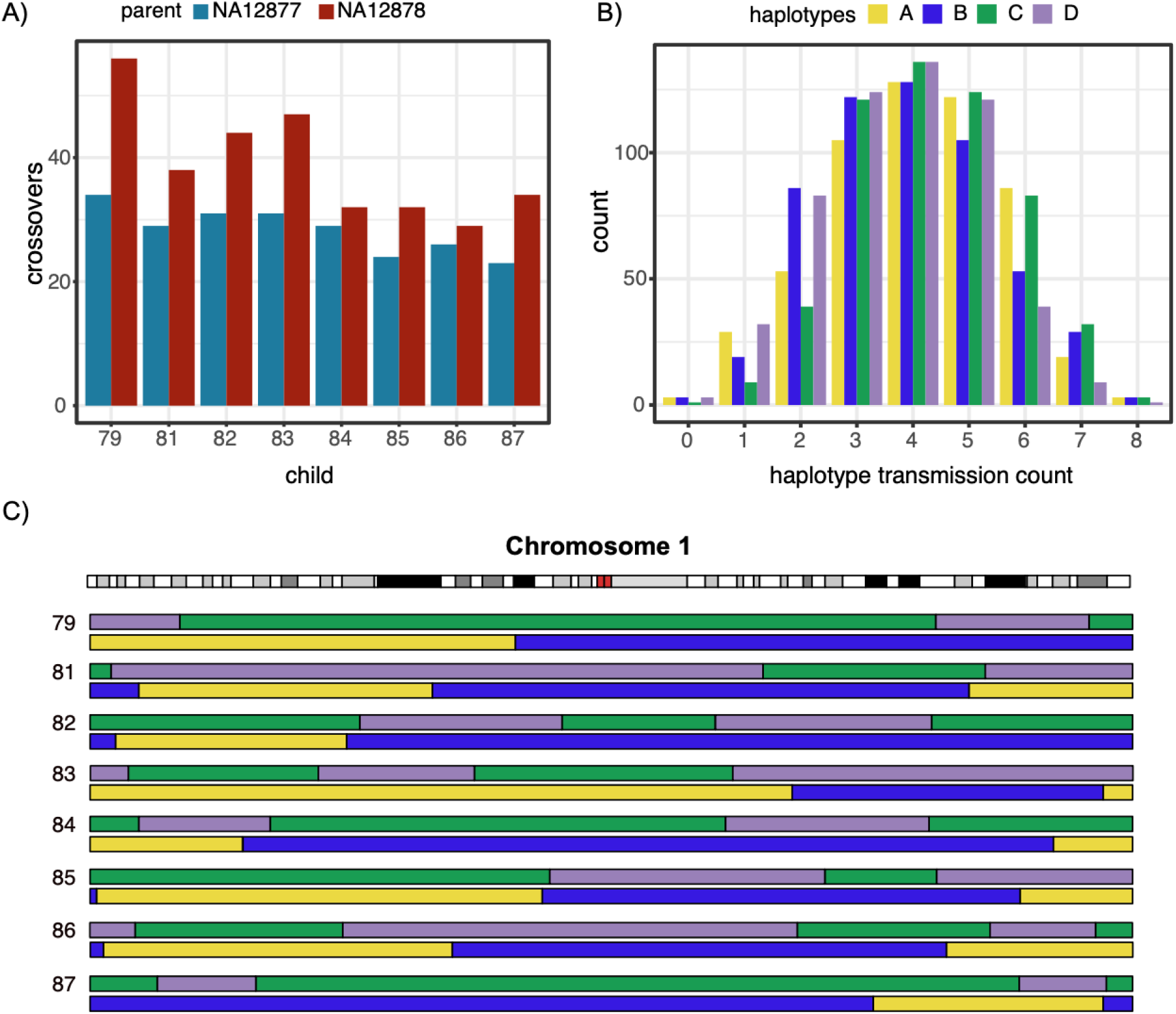
Tracking the four parental haplotypes. **A)** Recombination counts in the third generation stratified by the parent in which the recombination event occurred. **B)** Histogram of autosomal haplotype transmission counts. For example, zero means that one of the four haplotypes was not transmitted from G2 to G3. **C)** Recombination map for chromosome one. Haplotypes A and B (yellow and blue) are paternal and C and D (green and purple) are maternal. The prefix “NA128” has been removed from the sample IDs.

The haplotype inheritance map (inheritance vectors) constrains the genotype patterns possible at any given position and can be used to adjudicate all genomic variation because each variant must segregate congruently with the haplotypes (**Fig. 2C**). For this study, we use the terms true and accurate to describe variants whose genotypes in the members of this pedigree are consistent with the inheritance vectors. These true variants do not represent the full catalog of variants within this pedigree as some variants will be excluded because of genotyping errors, limitations of GRCh38, and or technological limitations. By combining multiple call sets and technologies, we increase the likelihood that a variant will be correctly genotyped across the pedigree and, thus, be included in our truth set.

### Small variants and confident regions analysis

We next identified “callable” regions of the genome where we expect to identify small variants if present in this family. To do so, we mapped ONT and PacBio high-fidelity (HiFi) data to GRCh38, establishing a union of technologies where all ten G2/G3 individuals have at least 10-fold coverage (**Fig. S4**). By combining two different technologies, we aim to minimize biases that may arise from differing read lengths or platform-specific sequencing bias. From this coverage-based map of confident regions, we excluded areas where pedigree-inconsistent variants are observed or where the reliability of alignment-based variant calling is uncertain (**Methods**). We also excluded the ten regions where all four haplotypes are not observed in the third generation. In total, the high-confidence callable regions span 2.77 Gb of the GRCh38 reference genome. The confidence regions will increase as pedigree-consistent variants are added back into the map.

We filtered and phased the small variant call set (SNVs and indels <50 bp) using our haplotype map, capturing ∼4.7-5.0M true SNVs and 384-900k true indels per technology (**Table 1, Table S4**). Approximately, 96% of the SNVs in each call set were considered true based on our consistency criteria (**Table S4**). Indels can be more difficult to genotype both accurately and consistently across the pedigree (Fang et al., 2014), and accordingly our indel truth rate ranged from 72-87% per call set. Of the variants that did not pass our pedigree inheritance test, 17-23% were identified in a single individual, likely representing false positives or *de novo* or somatic mutations (Porubsky et al., 2024) (**Table S4**). The remaining inconsistent variants had the same allele identified in multiple family members but failed the consistency check, indicating they are likely true variants with at least one genotyping error across the family.

**Table 1.** Counts of pedigree-consistent SNVs and indels within high-confidence regions.

| Pedigree Consistent |  |  | SNVs | Indels |
| --- | --- | --- | --- | --- |
| Illumina | PacBio | Nanopore |  |  |
| Yes | Yes | Yes | 4,290,226 | 317,997 |
| Yes | Yes | No | 317,872 | 308,642 |
| No | Yes | Yes | 189,834 | 24,409 |
| No | Yes | No | 63,027 | 89,094 |
| Yes | No | Yes | 25,262 | 8,096 |
| Yes | No | No | 42,214 | 205,143 |
| No | No | Yes | 66,128 | 11,608 |
| Total |  |  | 4,994,563 | 964,989 |

To create our consensus small variant benchmark, we normalized and merged these technology-specific truth sets (**Methods**). Importantly, to create a comprehensive truth set that benefits from the complementary strengths of different technologies, we included all variants if they were identified as pedigree consistent by at least one of the technologies. Variant merging resulted in 4,994,563 SNVs and 964,989 indels (**Table 1, Fig. S5**) For the SNVs, 96.6% were identified by at least two technologies and 3.4% were identified by a single technology (single technology breakdown: 25% Illumina, 36% HiFi, and 39% ONT). For the indels, 68% were identified by at least two technologies and 32% were identified by a single technology (67% by Illumina, 29% by HiFi, and 4% by ONT). It should be noted that based on our coverage analysis described above, we excluded 341,105 sites that were identified as pedigree consistent. Of the sites outside our high-confidence regions, 46%, 75%, and 44% had Illumina, HiFi, and ONT support, respectively.

There were 33,572 SNV and 59,493 indel loci that were excluded from our final truth set because they had different genotypes or conflicting alleles between technologies. We used the number of consistent and inconsistent sites to estimate an average discrepancy rate of 0.02% and 2.08% for the SNV and indel catalogs, respectively (**Methods**). The discrepancy rate corresponds to the probability that a pedigree-consistent site may be inconsistent between truth sets. These differences may be due to genotyping errors in some of the samples or nuances of the different aligners and variant callers. In particular, for the indels we expect this number to be an overestimate of the true error rate, as many of the 59,493 indel loci that were removed are not due to errors but rather different representations of the same variants. These discrepancies can often occur in the alignment and variant calling steps of different secondary analysis tools.

We sought to understand how variant calling and bioinformatic workflow discrepancies could lead to an underestimate of technology overlap. In other words, why are some variants in the truth set (14% of SNVs and 67% of indels) not supported by all three technologies? To answer this question, we examined the read-level evidence for all SNVs using a read-pileup-based analysis (**Methods, Fig. S6A-B**). At sites missing in one or more of the technology truth sets, we find 33.2%, 93.8%, and 98.3% had read-based evidence of an alternate allele, for Illumina, HiFi, and ONT, respectively. When missing an SNV call, Illumina data had the fewest sites with read support, which is expected since repetitive genomic regions require longer reads for confident mapping. The short reads had a lower average median map quality score (10 mapQ) compared to the long reads (60 mapQ)(**Fig. S6C**), and the short reads could not be reliably mapped at missing SNV sites. These results suggest that variant calling improvements could rescue variants missed by long-read sequencing technologies, but not those missed by short-read sequencing technologies.

### Comparison against other small variant benchmarks

Previous studies built benchmarks for this pedigree (Eberle et al., 2017) or NA12878 (Zook et al., 2016) primarily using short reads. Therefore, we expect that by using a combination of long- and short-read datasets across the pedigree, this study will be able to assess more of the genome. To confirm this, we compared our confident regions against those from previous studies. Our 2.77 Gb high-confidence region contains (190-260 Mb) more than either previous study: 2.51 Gb for GIAB 4.2.1, which excluded Chromosome X, and 2.58 Gb for the PG. Our high-confidence regions include 201 Mbp not present in either previous study, equivalent to the length of Chromosome 3, including difficult to analyze regions such as: (a) repeats (188 Mb), (b) low mappability regions (89 Mb), (c) segmental duplications (SDs; 42 Mb), and (d) coding sequence from parts of 2,833 protein-coding genes (13.9 Mb). Our truth set contains 218,077 SNVs and 185,558 indels within these difficult regions that are absent from both previous studies.

We next did a three-way comparison of our NA12878 SNV and indel truth sets against PG and GIAB (using BCFtools). There is a high concordance between the three truth sets with 76% (3.6M) SNVs and indels occurring in all three call sets (**Fig. S7**). There are 372,461 (8%) sites (171,446 SNVs and 201,015 indels) that are unique to the Platinum Pedigree dataset. Conversely, there are 235,345 (5%) sites that were not included in our dataset; of these sites 58% are outside our high-confidence regions. To estimate the true positive rate for Platinum Pedigree-only variants, we randomly sampled and manually inspected 100 SNVs and indels unique to the Platinum Pedigree truth set (**<u>Methods</u>, Table S5**). These variants are not found in either the PG or GIAB truth sets. We estimated the midpoint precision (between two reviewers) for the Platinum Pedigree-only SNVs and indels to be 50.5% and 79.5%, respectively.

Where the GIAB seeks to achieve reliable identification of errors (Olson et al., 2023) within defined benchmark regions at the expense of completeness, this study and the PG were designed to identify a more complete set of variants at the expense of specificity. To better understand the conflicting variants between Platinum Pedigree (PP) and GIAB, two independent reviewers scrutinized alignment data using Integrative Genomics Viewer (IGV), focusing on sites where one of the truth sets was called as homozygous reference. There are 4,658 conflicting positions (**Table 2**) of which 200 SNVs and 200 indels were manually reviewed (**Methods, Table S5**). Based on manual review, we note areas that may require further work including SNVs and indel calling proximal to structural variation, homopolymer indel accuracy, duplicate variants on the same haplotype, and cases where local haplotypes are missing a subset of variants due to pedigree filtering. By including more difficult regions it becomes more subjective to manually adjudicate complex variation but the conflicting variants in these regions only represent ∼0.1% of the truth set.

**Table 2.** Comparison between SNV and indel benchmarks in NA12878. Comparisons are based on hap.py analysis (subset of **Table S6**). False positives represent discrepancies between the truth and test call sets that are unique to the Test (Platinum Pedigree [PP]) call set and false negatives are unique to the truth call sets.

| Test | Truth | True Positive | False Positive | False Negative | Outside Truth regions | Outside Test regions | Genotype or allele conflict |
| --- | --- | --- | --- | --- | --- | --- | --- |
| PP | GIAB | 3,625,204 | 3,190 | 1,468 | 740,849 | 268,113 | 627 |
| PP | PG | 4,030,467 | 14,142 | 8,962 | 465,027 | 129,584 | 1,254 |

### Structural Variant (SV) analysis

The long reads also allow us to create a pedigree-validated truth set of SVs. Compared to SNVs, SVs are often represented differently causing complications in assessing and merging different call sets. To minimize these complications, we limited our SV analysis to the long-read datasets using both alignment- and assembly-based methods (**Methods**). This integrated call set consists of 35,662 SVs totaling 30.9 Mb of inserted, deleted, and inverted sequence. The majority (74.9%) of our integrated variants are supported by more than one call set, on average 2.6 different methods, and 25.1% of the calls are unique to only one method. Deletions are generally considered easier to identify and genotype than insertions, which is reflected in our results where 46.7% of the deletions are supported by all methods versus 22.6% for the insertions.

The difficulty of joint genotyping increases with the size of genetic variation, progressing from SNVs to indels to SVs. For example, small variant callers exhibit high pedigree concordance (∼95%), compared to the wide range observed amongst SV methods (41-86%) (**Table S7**). While assembly-based call sets include nearly twice as many variants as alignment-based methods, the number of variants that pass the inheritance genotype filters remains fairly consistent between both approaches (**Table S7**). For SVs that were pedigree consistent across multiple callers, 87.5% of deletions were consistent across all callers compared to 78.2% of insertions. The lower pedigree consistency for SVs (41-86%) compared to SNVs and indels highlights the need for better computational methods for more accurate and consistent SV genotyping and representation.

We next examined the distribution of SVs, sequence context, and reference bias in our truth set. As expected, we observe nearly fivefold more SVs (in binned counts) at chromosome ends and a higher frequency of insertions and deletions of 300 bp and 6 kb size (**Fig. 3A**), characteristic of Alu and LINE-1 elements, respectively (Audano et al., 2019). The majority (89%) of detected SVs falls within non-SD regions of the genome, while only 10.6% overlap SD regions. Nevertheless, SD-associated SVs affect four times more bases than those from non-SD regions despite being ten times less frequent (**Fig. 3B**). Detection accuracy can further be improved by utilizing a more complete and accurate reference genome, such as T2T-CHM13. This improvement is most evident in regions containing human satellites (including centromeric higher order repeats), where we now report 4,048 pedigree-consistent SVs affecting a similar number of bases as SD-associated variants (**Fig. S8A-B**). The main SV call set is relative to GRCh38, where we see an increased overall number of insertions (n=9,641) in comparison to the T2T-CHM13 reference (n=6,782), which is reflected in an increased insertion to deletion ratio. This bias is not observed for deletions, where we see nearly equal numbers (∼6,200) across both references (**Fig. S8C, Fig. S9**).

**Fig. 3.**
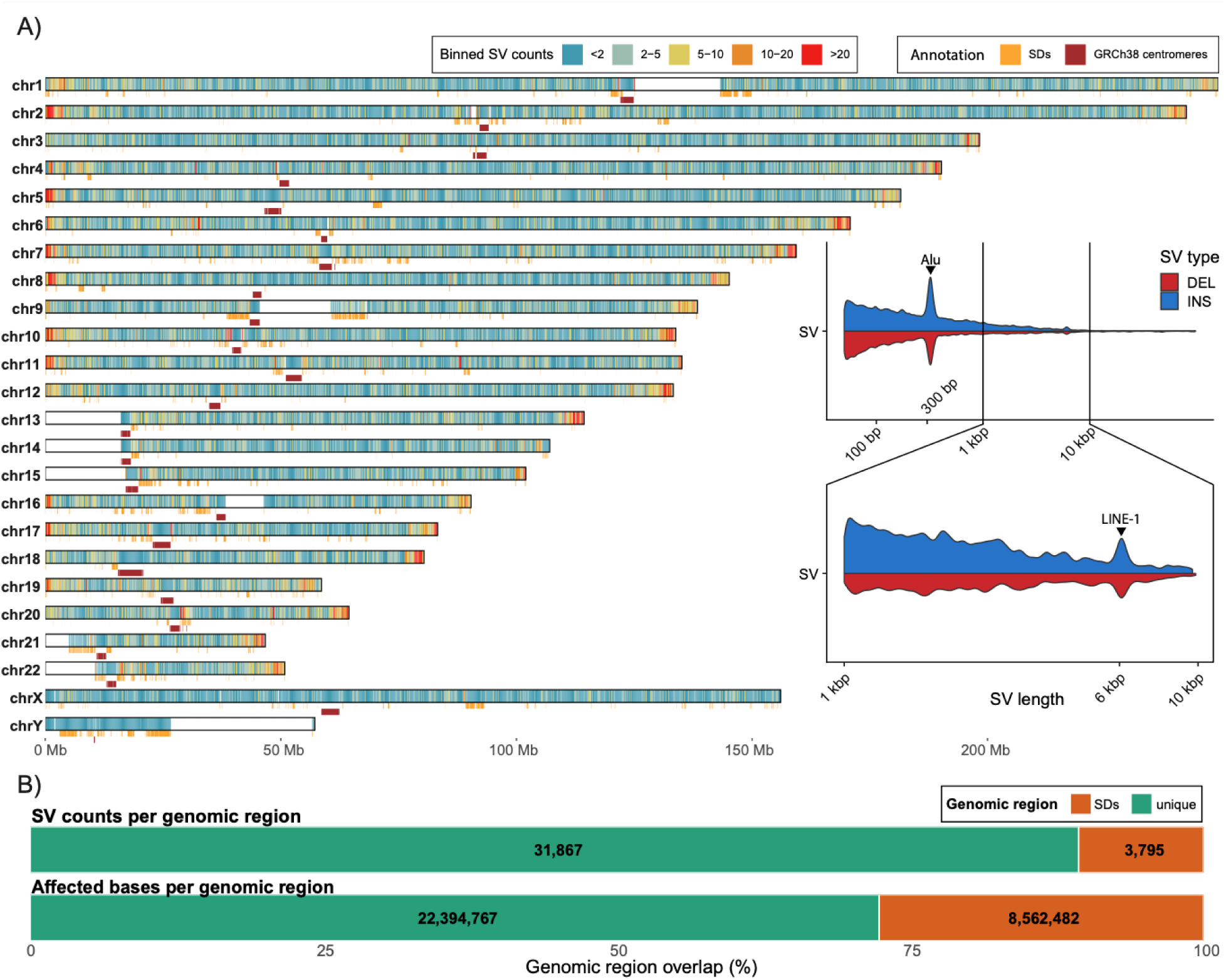
Structural variant (SV) density across the pedigree relative to GRCh38. **A**) Density of SVs with respect to GRCh38. SVs are counted in 200 kb long bins. GRCh38 SD annotation is shown as orange-colored rectangles below each chromosome along with the positions of centromeres shown as red rectangles. Inset shows a size distribution of SVs for insertions (blue) and deletions (red) above and below the midline, respectively. We mark increased frequency of ALU elements around 300 bp as well as LINE-1 elements around 6 kbp. **B**) Top bar shows counts of SVs overlapping non-SD (green) and SD (orange) space. Bottom bar shows the amount of affected base pairs by SVs overlapping non-SD (green) and SD (orange) space.

Our merged SV call set includes 24,160 SVs in NA12877 and 24,171 SVs in NA12878 (**Table S8**). Prior to this study, a comprehensive SV truth set had not been created for this pedigree, although NA12878 was included in two studies that used either short reads or lower-depth long-read data to estimate SV counts in the population. The first study using short-read sequencing identified just 3,229 SVs in NA12878 (Sudmant et al., 2015). A second study, using relatively low depth (16.9-fold) ONT long-read data, identified 19,029 SVs in NA12878 (Schloissnig et al., 2024). Even at lower depths, long reads are able to confidently identify more SVs per sample than short reads, although our study, with an average depth of 52-fold coverage, identifies and validates 26% more SVs in this sample. A more recent study using a diverse set of samples using high-depth (37-fold) long reads found an average of 24,543 SVs per individual (Gustafson et al., 2024), similar to our results, though the samples included in that study do not overlap this family.

### Tandem repeat (TR) analysis

TRs are among the most difficult and mutable variants (Depienne & Mandel, 2021) to profile. Here, we genotyped 7,722,730 tandem repeats (Porubsky et al., 2024) across the CEPH-1463 pedigree samples with a recently developed Tandem Repeat Genotyping Tool, TRGT (Dolzhenko et al., 2024). Since the vast majority of these loci are short, non-polymorphic repeats that can be accurately resolved by generic small variant callers, we focused on a subset of 650,610 TRs where: (a) at least two distinct alleles are observed, (b) the mean allele purity score is at least 0.5 (**Methods**), (c) each allele is observed at least twice (to exclude *de novo* mutations and other errors), (d) all alleles are present and have at least one motif copy identified within them, and (e) at least one allele spans ≥10 bp (**Table S9**). Analysis of concordance between inheritance vectors and TR allele sequences revealed that 82.62% (N=537,540) of these TRs are pedigree consistent. Consistency rates were similar across allele lengths, motif lengths, and allele purities (**Fig. S10A-C**), with the exception of homopolymer repeats (**Fig. S10B**) where the consistency rate was lower (71.7%).

Unlike indels and SVs, which are defined based on length, TRs are characterized by the repetitive nature of the sequence that comprises these events (**Fig. 4**). Thus, changes in repeat length quantified in this TR call set may overlap our catalogs of insertions and deletions, which are defined by size (e.g., indels and SVs). TRGT reports the full length of the repeat region rather than changes relative to the reference. We found that 11,347 (31.8%) of the SVs and 275,307 (30%) of the indels in our call sets are contained within our pedigree-consistent TR truth data. Excluding overlapping indels and SVs, our merged truth data includes 4,719,711 SNVs, 767,795 indels, 537,486 TRs, and 24,315 SVs (**Table 3**).

**Fig. 4.**
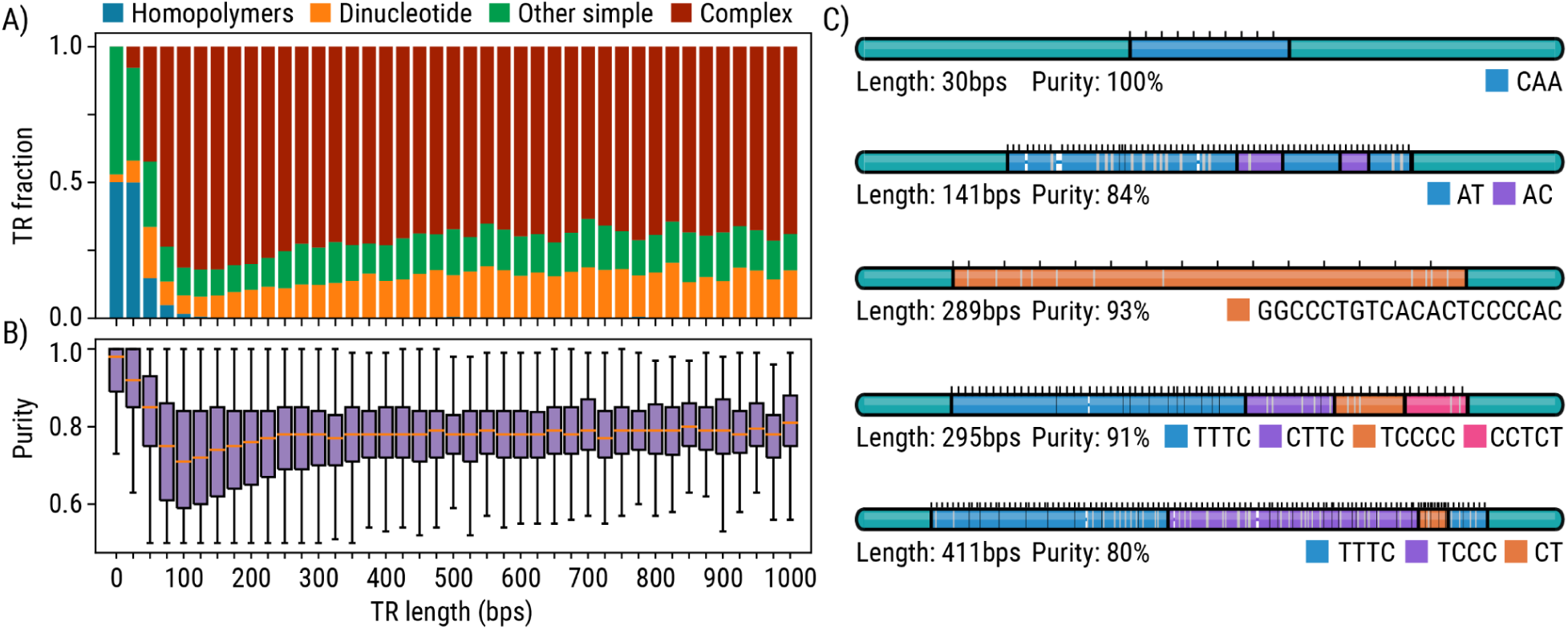
Repeat content characteristics. **A)** Proportions of homopolymers, dinucleotide TRs, simple TRs with longer motifs, and complex TRs containing multiple motifs stratified by the mean allele length. **B)** Allele purities stratified by the mean allele length. **C)** Example TR alleles of different sizes, purities, and composition.

**Table 3.** Summary of variation in CEPH-1463. The average caller overlap is calculated as the mean number of methods that support each call. The denominator for average caller support is three for SNVs and indels, one for tandem repeats, and four for SVs. All variant counts are non overlapping between SNVs, tandem repeats, and SVs. SVs, SNVs, and indels that overlap TR-defined regions were removed.

|  | Count | Total Len. (bp) | Avg. Len. (bp) | Average caller overlap |
| --- | --- | --- | --- | --- |
| SNVs | 4,719,711 | 4,719,711 | 1 | 2.8 (94%) |
| Indels | 767,795 | 2,890,759 | 3.3 | 1.96 (66%) |
| Tandem repeats | 537,540 | 2,119,755 | 5.6 | 1 (100%) |
| Deletions | 13,460 | 14,040,611 | 1,531 | 2.9 (73%) |
| Insertions | 15,089 | 13,749,758 | 911 | 2.3 (60%) |
| Inversions | 55 | 365,263 | 6,641 | 1.1 (33%) |
| Total | 6,053,650 | 37,885,857 | 6.25 | N/A |

### Utility for methods development

The GIAB datasets (including NA12878) have been used to train DeepVariant (Wagner et al., 2022) in the past. To demonstrate how the more extensive Platinum Pedigree truth set would further improve variant calling, we modified the DeepVariant training labels to incorporate the Platinum Pedigree truth dataset for NA12878 and compared performance of a model trained on the Platinum Pedigree labels with the prior release of DeepVariant (v1.6), which was trained using only the GIAB labels. We evaluated a 35-fold HiFi sample on NA12878, comparing accuracy on Chromosome 20, which is never used for training. On the GIAB benchmark set (v4.2.1), a model trained with the Platinum Pedigree NA12878 labels had 8% fewer errors (**Table S10**), driven by roughly equal improvements in precision and recall in both SNPs and indels. On the more extensive Platinum Pedigree truth set, a model trained with Platinum Pedigree labels had 34% fewer errors (**Table S11**) driven mostly by improvements in SNV recall (0.9613 to 0.9783), indel recall (0.9463 to 0.9556), and indel precision (0.9678 to 0.9752). The difference in the number of errors (a hundred for GIAB and thousands for Platinum Pedigree) reflects that the Platinum Pedigree truth set covers more of the genome, including the more difficult to analyze parts of the genome.

## Discussion

In this study, we leveraged sequence data from three sequencing platforms (Illumina 2x150bp, PacBio HiFi, and ONT-UL) to create a high-confidence truth dataset for the CEPH-1463 pedigree, encompassing two generations of eight siblings and their parents. The dataset spans 2.77 Gb of the GRCh38 genome, capturing a wide range of variant types, including a nonoverlapping set of 4,719,711 SNVs, 767,795 indels, 537,486 TRs, and 24,315 SVs. This resource enables precise delineation of genetic variation across diverse variant classes.

Compared to the GIAB benchmark set (v4.2.1), our dataset identifies 11.6% more SNVs and 39.8% more indels in NA12878. This improvement is driven by the use of newer sequencing technologies and the application of pedigree-based validation, allowing variant accuracy assessments without filtering for sequence context. While GIAB is designed to minimize false positives, our broader dataset captures variants in more challenging genomic regions, facilitating the evaluation of sequencing technologies and the enhancement of variant detection methods. For example, the expanded SNV and indel dataset was used to retrain DeepVariant, reducing error rates by up to 34%, with these gains being integrated into DeepVariant v1.8.

The stringent filtering pedigree-consistency criteria means that we will filter some true variants due to genotyping errors in as few as a single sample. As variant callers improve, many of these excluded variants may be recovered. In particular, our analysis demonstrated challenges in detecting and genotyping complex variants like SVs and TRs, where up to 60% and 42%, respectively, fail inheritance validation, despite strong supporting evidence in multiple individuals. These failed variants will highlight areas where algorithmic improvements are needed. In addition to validating variants that are correctly genotyped across the pedigree, the inheritance vectors can also be used to pinpoint samples with potential genotyping errors, helping refine variant calling algorithms.

A limitation of this study is that many conflicts between call sets were not resolved. These conflicts were most likely to occur in difficult-to-interpret regions where there are clusters of variation of repetitive sequence (e.g., low-complexity repeats). Many times these inconsistencies occur because different aligners and variant callers can represent variation in different, and apparently conflicting, ways. This was clear when comparing indels between different truth sets. Future work will test pedigree assessment of localized *de novo* assemblies and additional technologies as a way to resolve the larger scale sequence of these regions.

This study relied completely on alignment-based variant calling but for highly variable and complex regions of the genome we will extend this work to include the diploid assemblies as a way to assess the quality of these assembled haplotypes. For example, assemblies are prone to errors in very long repeat regions like the KIV-2 repeat (one of the longest human VNTRs) and regions of duplicated sequence that are prone to frequent gene conversion and copy number variation like the complicated region where *SMN1* and *SMN2* are located. For these types of regions, combining the assemblies with the inheritance map will allow us to develop a first-level assessment of the accuracy of the genome assemblies. Ongoing work will apply our pedigree-based filtering framework to phased genome assemblies to assess the accuracy of the assemblies in these complex regions.

The variant benchmark presented here is not meant to represent a complete truth set but instead represents a starting point that will continually be improved as new variant callers are developed. As this benchmark is improved to more accurately represent the full diploid genomes of this family, we expect that the results will continue to help tool developers and train new and ever-improving variant callers to resolve variability in more of the dark genome. Beyond the utility for developers, it can be increasingly difficult to validate the performance of sequencing pipelines to accurately genotype difficult variants, such as SVs and TRs. This benchmark will enable labs to validate the performance of variant calling for SVs and TRs so that they can be implemented in clinical settings.

## Acknowledgments

We would like to acknowledge Daniel Baker for exploring SV merging methods and Tonia Brown for editing the manuscript. This work was supported, in part, by US National Institutes of Health (NIH) grants HG010169 to E.E.E. E.E.E. is an investigator of the Howard Hughes Medical Institute. Certain commercial equipment, instruments, or materials are identified to specify adequately experimental conditions or reported results. Such identification does not imply recommendation or endorsement by the National Institute of Standards and Technology, nor does it imply that the equipment, instruments, or materials identified are necessarily the best available for the purpose.

## Contributions

Z.K., C.N, D.P. T.M., W.J.R., S.L., E.D., A.C., J.H., C.S., E.E.E. and M.A.E. wrote the manuscript. Z.K., C.N, D.P. T.M., W.J.R., S.L., E.D., N.K., W.T.H., A.C., J.H., C.S., P.C., S.M., A.C, M.D.O., J.M.Z and K.C. processed data and did analyses. K.M., K.H. W.S.W., C.F., and C.L. generated sequencing data. Z.K, D.P., A.C, H.D., J.M.Z, P.M.L., E.G., J.S., P.M.L, L.B.J, A.R.Q, E.E.E, and M.A.E. provided oversight and designed experiments.

## Methods

### Data generation

DNA was extracted from whole blood and sequenced using Illumina, PacBio HiFi, and ONT-UL instruments. Strand-seq data was generated from cell lines as described in [Porubsky et. al. 2024]. For this study, we aligned the Illumina, PacBio, and ONT data to GRCh38 to generate small variant calls (SNVs and indels). SV calls were made using the long-read data (PacBio and ONT) using both alignment-based genotypers and assembly-based callers. The Strand-seq data was used to confirm the haplotype transmission calculated for this pedigree (see below).

### Ethics declarations

Human subjects: Informed consent was obtained from the CEPH/Utah individuals, and the University of Utah Institutional Review Board approved the study (University of Utah IRB reference IRB_00065564).

### Data availability

For the open consent samples, the data are released on Amazon Open Data. The Amazon S3 bucket contains the sequencing data, assemblies, variant calls, and additional files documented here:

https://github.com/Platinum-Pedigree-Consortium/Platinum-Pedigree-Datasets

Samples with controlled access can be found at dbGaP under the accession **XXXX (in progress**).

### Inheritance vectors

A hidden Markov model was developed where, at a heterozygous SNV locus, the hidden state was the true inheritance of the paternal/maternal haplotypes, and the observed state was the presence/absence of an SNV call in each of the G3 children. This enabled us to identify the most probable sequence of inheritance vectors across a chromosome using the Viterbi algorithm. The transition matrix defines the probability of a transition from any one inheritance state to another (i.e., recombination). The emission matrix defines the probability of a set of child calls representing a given inheritance state. The matrices were defined according to the following equations, when *i* ≄ *j*:

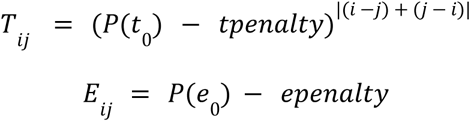

Where *|(i - j) + (j - i)|* is the number of differences between the *i* and *j* observed states (sets of inheriting children), *P(t_0_)* the initial transmission probability*, tpenalty* the transmission difference penalty, *P(e_0_)* the initial emission probability, and *epenalty* the emission difference penalty. Parameters *P(t_0_), tpenalty*, *P(e_0_)*, and *epenalty* were iteratively refined to recapitulate previous vectors and correctly identify missing recombinations. Programmatically identified inheritance vectors were compared to an initial sketch of inheritance vectors identified by applying a depth filter (>1.67 or <0.27 times the sample mean, equivalent of a 50-fold upper bound and 8-fold lower bound for a 30-fold depth sample) to SNV calls from DeepVariant. After depth filtering we identified runs of greater than ten SNVs supporting the same inheritance from NA12877 and NA12878, respectively. These were manually refined by removing unlikely changes in inheritance (i.e., close range double recombinations or gene conversions), closing gaps between vectors using unfiltered calls and adding vectors between blocks separated by more than one recombination where SNV support existed for the intervening inheritance pattern. Missing recombinations were identified by observing blocks of pedigree-violating variants occurring in the same child, which matched the location of recombinations identified via assembly-based caller PAV and Strand-seq. The Viterbi approach identified an additional 20 inheritance blocks in GRCh38, for a total of 539 recombinations and 560 inheritance blocks. Updated inheritance vectors resulted in a decrease in single-pedigree violations (147,603 to 60,394) and an overall increase in pedigree-consistent small variants from HiFi (4,988,878 to 5,068,085). The resulting haplotype inheritance blocks covered 97.4% of the GRCh38 autosomes and Chromosome X. These final vectors were compared with results from Strand-seq reported in **Table S2**.

### Small variant analysis

Three variant callers were used to generate a maximally sensitive and pedigree-consistent call set (**Table 1**). Joint variant calling was carried out across the pedigree, and then the second and third generations were isolated from the broader call sets. Small variant counts are reported in **Table S3**. After variant calling, small variants were filtered (QUAL ≥ 20) and normalized. Variant sites with more than four alleles or sizes >49 bp were excluded. Normalization was applied to standardize alleles. The command applied is listed below:

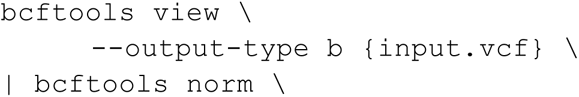

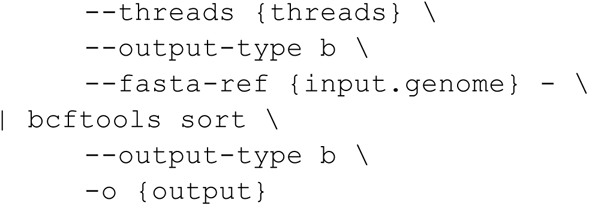

One inherent issue of merging distinct variant calls is overlapping variants. For example, a deletion cannot span an SNV on the same haplotype. Similarly, insertions at the same position and haplotype are invalid. In the two sets of parental haplotypes (NA12878 and NA12877) there were 93,065 overlapping variants. We removed these variants (limited only to that site) from the truth set as the site is ambiguous. To evaluate platform-specific variants, we used `minipileup` (https://github.com/lh3/minipileup) to obtain allele counts in the G2 generation from uniquely aligned reads (MAPQ > 0) across the three sequencing platforms.

### DeepVariant calling

The human-WGS-WDL software pipeline was used to call small variants from HiFi data (https://github.com/PacificBiosciences/HiFi-human-WGS-WDL/releases/tag/v1.0.3). The pipeline aligns, phases, and calls small variants with DeepVariant. GVCF files were merged with GLnexus. Software versions used are documented (https://github.com/PacificBiosciences/HiFi-human-WGS-WDL/tree/v1.0.3?tab=readme-ov-file#tool-versions-and-docker-images).

### ONT clair3 calling

Clair3 (v1.0.7) variant calls were made based on the alignments with default models for ONT (ont_guppy5) data with phasing and gVCF generation enabled. Variant calling was conducted on each chromosome individually and concatenated into one VCF. gVCFs were then fed into GLNexus with a custom configuration file.

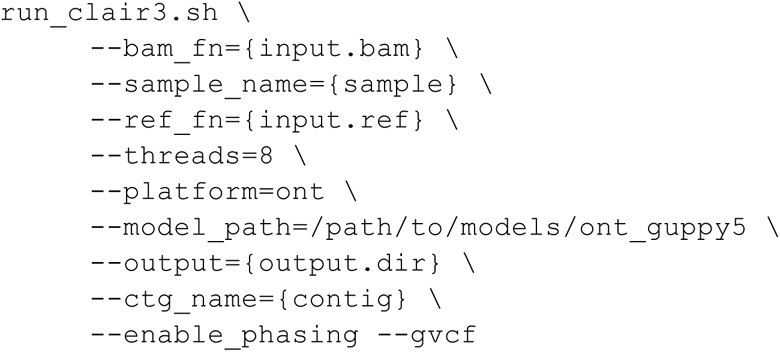

### DRAGEN calling

DRAGEN v4.2.4 was used to align and call variants from Illumina short-read data against GRCh38 (hg38-alt_masked.cnv.graph.hla.rna-9-r3.0-1) and the T2T (chm13v2.0_maskedY_rCRS) reference. References can be found at: https://support.illumina.com/sequencing/sequencing_software/dragen-bio-it-platform/product_files.html

### Pedigree filtering

Code was implemented to filter and phase variants based on the second-to-third generation haplotype transmissions, using the logic outlined by Eberle 2017 (https://github.com/Platinum-Pedigree-Consortium/Platinum-Pedigree-Inheritance/blob/main/code/rust/src/bin/concordance.rs). The concordance program accepts a VCF (as detailed in small variant analysis) and the inheritance vectors (as detailed in defining inheritance vectors). The concordance code generates all possible combinations of valid genotypes over a haplotype (region) and tests if the observed genotypes at a site match a valid combination. Sites that pass the inheritance filtering are kept if they contain a genotype call for every individual (excluding any site with no calls). Using the inheritance vectors we can directly phase genotypes for bi-allelic and multi-allelic sites. There is a bijective mapping of observed genotypes in the VCF with the haplotype structure with the exception of all heterozygous genotypes, where the phase is ambiguous. We exclude sites where all individuals are heterozygous from the truth set. The command used to filter and phase the variants is:

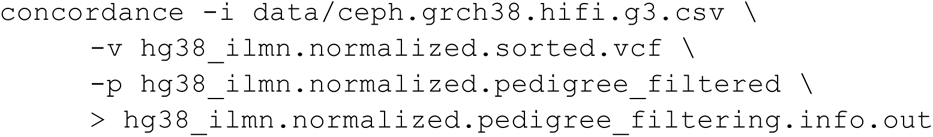

Each pedigree-consistent variant call, encoded as a VCF record, was reduced to a concatenated string consisting of chromosome, position, ref, alt, genotypes, and caller. The reduced representation of the calls was then concatenated across callers, sorted, and merged. Only identical records were merged, and sites where the genotypes differ between callers were excluded from the final truth set. VCF records were reconstituted from the merged call sets. Each VCF record reports which technologies and callers support the site.

The process of building a small variant truth set for NA12878 is encapsulated in a Snakemake: https://github.com/Platinum-Pedigree-Consortium/Platinum-Pedigree-Inheritance/tree/main/pipelines/smallvar-filtering

### Calculating error rate of Platinum Pedigree truth set

To estimate an error rate we enumerate conflicting calls observed early in the variant merging process. The simplest error is when two variants share the same site and same alternate alleles, where one of the ten genotypes mismatch between two call sets. The next error is containment, when one variant represents one fewer allele. Take, for example, two alternate alleles sets `A,ATT` and `A`. The `A` allele is contained in the other call, yet missing one allele. In this case, we keep the variant with more alternate alleles and mark the other as an error. The last type of error is when there are two variants on the same haplotype but overlap. For example, an SNV called within a deletion on the same haplotype, which is not biologically possible.

| Error category | #SNVs | #indels |
| --- | --- | --- |
| Genotype mismatch | 3,809 | 1,125 |
| Allele mismatch | 1,134 | 8,563 |
| Overlapping alleles | 28,629 | 49,805 |
| Total | 33,572 | 59,493 |
| Estimated error rate | 0.003605508 | 0.05918883 |
| Single tech support rate | 0.05993709 | 0.3528733 |
| Single tech estimated error | 0.0002161037 | 0.02088616 |

### Calculation of the error rates, based on the counts of shared variants between call sets

The numerator is the total number of mismatches between technologies, including genotype mismatching, allele mismatches, and overlapping alleles. The denominator is the number of times two and three technologies share a call. For example, there are 567k times two technologies support the same call plus 4.3M sites where all three technologies support the same call. The number of calls where all three technologies support the same event generates two sets of comparisons and therefore is doubled.

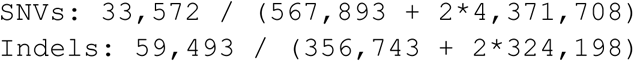

### Manual inspection of Platinum Pedigree unique small variants

To determine the true positive rate of Platinum Pedigree-specific small variants, `bcftools isec` was used to compare the Platinum Pedigree truth set with the Platinum Genome and GIAB truth sets to isolate NA12878 variants unique to the Platinum Pedigree truth set. Afterwards, 100 biallelic SNVs and indels were randomly selected for manual inspection using IGV. A variant was deemed a true positive if the variant was unambiguously present in the source sequencing platform with the expected variant allele fraction, given the genotype (i.e., 30%-70% for heterozygous variants and 80%-100% for homozygous variants). Results of the manual inspection are summarized in **Table S5** and IGV screenshots are available as part of Supplementary Information.

### Read support of single-nucleotide variants

The reference and alternate allele counts, along with the MAPQ scores at each variant site, were obtained for all biallelic SNVs (∼4.93M sites) in the Platinum Pedigree truth set for the G2 generation members (NA12877 and NA12878) across the three sequencing platforms using `samtools mpileup`. If either parent had at least three reads supporting the minimum alternate allele count, we considered there to be read-based evidence for the alternate allele in the sequencing platform of interest.

### Manual inspection of conflicts between Platinum Pedigree and GIAB truth sets

To identify conflicts between Platinum Pedigree and GIAB truth sets, we used hap.py with GIAB 4.2.1 benchmark as the truth set and Platinum Pedigree as the query:

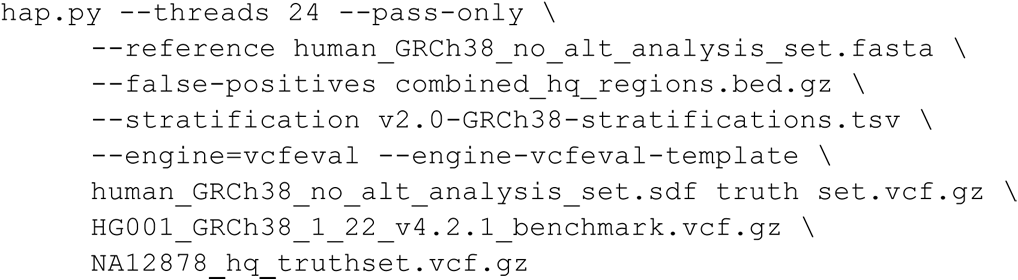

Afterwards, 100 false positive SNVs and indels and 100 false negative SNVs and indels were randomly selected for manual inspection using IGV (**Table S5**). We scrutinized the evidence for the alternate allele specific to PP and GIAB truth set in the Illumina, HiFi, and ONT data. We required that the variant is biallelic, variant representation did not change during the comparison, and GIAB is homozygous reference for false positives and Platinum Pedigree is homozygous reference for false negatives. We ignored genotyped sites where the same variant was called in both truth sets but had different genotypes for manual inspection (e.g., the variant was called a homozygous alternative in the GIAB benchmark, while it was identified as a heterozygous variant in the Platinum Pedigree truth set).

### High-confidence regions

To collate callable regions across the genome where we are confident that the samples are homozygous for the reference allele, we next identified places where, based on alignments, we would have identified small variants if they are present in this family. To do so, ONT and HiFi data were mapped to GRCh38 establishing the union of technologies where all ten G2/G3 individuals have at least 10-fold coverage in either technology (**Fig. S4**). By taking the union of two different technologies we avoid biases that arise from differing read lengths or platform-specific sequencing bias. From this coverage-based map of confident regions, we removed places where variants are observed or we cannot be confident of the performance of alignment-based variant calling. Positions where either long-read technology (ONT and PacBio HiFi) confidently maps (MAPQ ≥ 10) and every individual has at least 10-fold coverage are the starting high-confidence regions (https://github.com/Platinum-Pedigree-Consortium/Platinum-Pedigree-Inheritance/blob/main/analyses/hcr_regions.md). The union of the technologies yields 668 regions spanning 2.85 Gb of GRCh38. We intersect these regions with the haplotype map, as we can only establish confidence in regions where we have coverage and established inheritance vectors, resulting in 2.83 Gb of GRCh38 broken into 968 regions. There are 828 GRCh38 regions, including PARs, VDJ, and gaps that have been previously identified as problematic (Dwarshuis et al., 2023). We removed these regions resulting in 2.829 Gb across 1,082 regions. Next, we removed 40 Mb of the genome where one haplotype is missing from the inheritance vectors across all G3 samples (**Table S3; Fig. S3**). All variation was then removed (including structural variation, see following section) and only pedigree-consistent variants were added back into the high-confidence regions. In the end, there were 2.77 Gb of the genome deemed high confidence, broken into 1,054,558 regions.

### SV analysis and integration

SVs were detected by read-based calling and genome assembly-based calling. For read-based calling we used pbsv, sniffles (Smolka et al., 2024), and sawfish (Saunders et al., 2024) on the PacBio HiFi data. Two assembly calling tools were used: Phased Assembly Variant Caller (PAV) (Ebert et al., 2021) and PanGenome Graph Builder (PGGB) (Garrison et al., 2023). The assembly-based callers were run on aligned Verkko (Rautiainen et al., 2023) assembled contigs. Each call set was filtered to remove events less than 50 bp in size. The SV counts are listed in **Table S7** and range from 30-60k events. For the integrated call set we started with the most pedigree-consistent caller, sawfish, followed by sniffles (ONT), PGGB, and PAV. We decided to exclude pbsv and sniffles (HiFi) calls as they largely overlapped with sawfish. By using assembly-, ONT-, and HiFi-based SV calling we maximized SV discovery.

The sawfish joint-genotyping workflow was run over all ten samples in the second and third generations of the pedigree. Briefly, the workflow steps are (1) identification of SV signatures in each sample, (2) per-sample local assembly of SV haplotypes corresponding to each signature, (3) merging SV haplotypes across samples, and (4) per-sample genotyping of each merged SV haplotype by assessment of read support. Sawfish results are post-processed to select for passing variants at least 50 bp long and excluding breakpoints. The sawfish “HQ” set additionally removes variants with a genotype quality (‘GQ’) of at least 40 for every sample in generations G2 and G3.

To create the exclusion track for small variants, the sawfish joint sample VCF was filtered to remove indels less than 50 bp, and inversion calls were replaced with their component breakends. The variants were further down-selected to only retain calls with a non-reference genotype in NA12878 passing all filters. From the remaining variants, we defined the excluded region as any region that intersected within 25 bp of any remaining indel variants, or within 50 bp of any other remaining variants in the set.

SV normalization is performed prior to merging pedigree-consistent variant calls from different SV calling methods. This process standardizes SV representations in VCF files across different callers by removing method-specific information, unifying variant identifiers, and harmonizing common fields between the callers.

The SV merging strategy uses an interval tree-based approach to combine calls from multiple SV callers (sawfish, sniffles, PGGB, PAV). First, SVs from a designated primary caller (sawfish) are used to construct an initial interval tree. Each SV is represented as an interval in the tree, with a defined flank length of 200 bp to account for potential positional differences between callers. Variants from additional SV callers are then iteratively merged into this primary tree. For each new SV, there is a search for overlapping intervals in the existing tree. If an overlap is found, the new SV is added to the list of supporting variants for the existing SV entry. If no overlap exists, a new interval is introduced into the primary tree.

After merging, a selection process refines the supporting variants for each primary SV, addressing two scenarios. First, a supporting variant from a single caller may initially overlap with multiple primary variants. Second, multiple supporting variants from the same caller may overlap with a single primary variant. The method ensures that each supporting variant from any given caller ultimately supports only one primary variant. In instances where a supporting variant overlaps multiple primary variants, it is assigned to the most similar primary variant based on SV length similarity and parental genotype consistency. Conversely, when multiple supporting variants from the same caller overlap a single primary variant, the best supporting variant is selected among these candidates using the same criteria. Specifically, the criteria for determining the best match between supporting and primary variants involve comparing the length differences and genotype consistency. A length difference threshold of 30 bp is used to decide whether a supporting variant is sufficiently similar to a primary variant. If the length difference is within this threshold, the genotypes of the parents are compared. The genotypes must match between the primary and supporting variants, meaning both variants must show the same allelic configuration in the parents. Within the matching genotype-consistent variants, the variant with the smallest possible length difference is selected as the best match. This approach prevents over-counting of support, ensures each supporting variant contributes to only one merged SV call, and maintains a balanced representation of evidence from different callers.

### Structural variation comparison

To compare the integrated call set of SVs from different callers to the phase three 1000 Genomes Project (1KG) call set and our pedigree-consistent TR call set, a similar interval-tree-based strategy was used. All call sets were filtered to only include calls with alternate alleles from the NA12878 sample.

For the 1KG SV call set, only a subset of variants—specifically deletions—was included in the comparison. Each SV was represented as an interval in an interval tree, with a defined flanking length of 200 bp to accommodate positional discrepancies. SV intersections were determined by identifying overlapping intervals between the query interval and the search tree. Potential overlaps were further refined by calculating size similarities between the query and overlapping intervals based on SV length. A similarity threshold of 70% was used to determine if an overlap was significant. Notably, setting the threshold to 90% still resulted in an overlap of 83.8% of 1KG deletions in our merged call set, while a threshold of 95% yielded 80.2% overlap. Note that using sequence similarity was not feasible for this analysis, as many of the 1KG SV calls are symbolic, lacking sequence-level resolution.

In the comparison with TR calls, SV calls were no longer categorized by type. Furthermore, no flanking length was added to the intervals, as TRs are defined relative to predefined loci with fixed windows. Since TRs are characterized by established boundaries, adding flanking regions could introduce noise and lead to false positives by including unrelated nearby genomic features. For each SV interval, the nearest neighboring TR interval was identified using the corresponding TR interval tree. Containment was used to determine whether SVs were entirely within TR regions, ensuring that only complete overlaps were considered. Additional analysis, testing for interval intersection rather than strict containment, increased the number of TR-overlapping SVs by 2.58%.

### Filtering code

https://github.com/Platinum-Pedigree-Consortium/Platinum-Pedigree-Inheritance/blob/main/pipelines/sv/process.sn

### Merging code

https://github.com/Platinum-Pedigree-Consortium/Platinum-Pedigree-Inheritance/blob/main/analyses/StructuralVariant.md

### Tandem repeat analysis

Tandem repeats were genotyped with TRGT 1.1 (Dolzhenko et al., 2024). The allele purity scores were also calculated by TRGT. A purity score of 1.0 means that the corresponding repeat allele consists of the perfect copies of motifs specified in the repeat catalog file, while purity scores close to 0.0 are assigned to alleles that have little resemblance to the specified motifs. The Jupyter notebook capturing the described tandem repeat analysis is available https://github.com/Platinum-Pedigree-Consortium/Platinum-Pedigree-Inheritance/tree/main/pipelines/tandem-repeats).

### Retraining DeepVariant and running Platinum Pedigree models

Previous DeepVariant models were trained with GIAB labels for HG001-HG007 (excluding HG003 as a holdout). The training dataset breakdown by sample and training examples are detailed at: https://github.com/google/deepvariant/blob/r1.6.1/docs/deepvariant-details-training-data.md. We used the same training configuration for DeepVariant but replaced the truth label for HG001 with the Platinum Pedigree labels, resulting in a model trained on samples with labels from both Platinum Pedigree genomes and from GIAB. Chromosome 20 is not used during training and is used as a holdout to assess accuracy.

After generating the retrained model, we ran both the prior release version of DeepVariant (v1.6.1) and the new model, as described in the PacBio case study: https://github.com/google/deepvariant/blob/r1.6.1/docs/deepvariant-pacbio-model-case-study.md

After the evaluation demonstrated higher accuracy with the Platinum Pedigree-trained model, whether evaluated on the GIAB or Palladium truth, we released the new model for anyone to run.

The model files are available at:

https://storage.googleapis.com/brain-genomics-public/research/platinum/model/checkpoint-2035200-0.99245-1.data-00000-of-00001

https://storage.googleapis.com/brain-genomics-public/research/platinum/model/checkpoint-2035200-0.99245-1.index

https://storage.googleapis.com/brain-genomics-public/research/platinum/model/example_info.json

To run this DeepVariant model in the manner run for this study, use the v1.6.1 release of DeepVariant and execute the following command (assuming one has created a folder input and output and placed BAM and fa.gz, fa.gz.fai, and fa.gz.gzi files in the input directory):

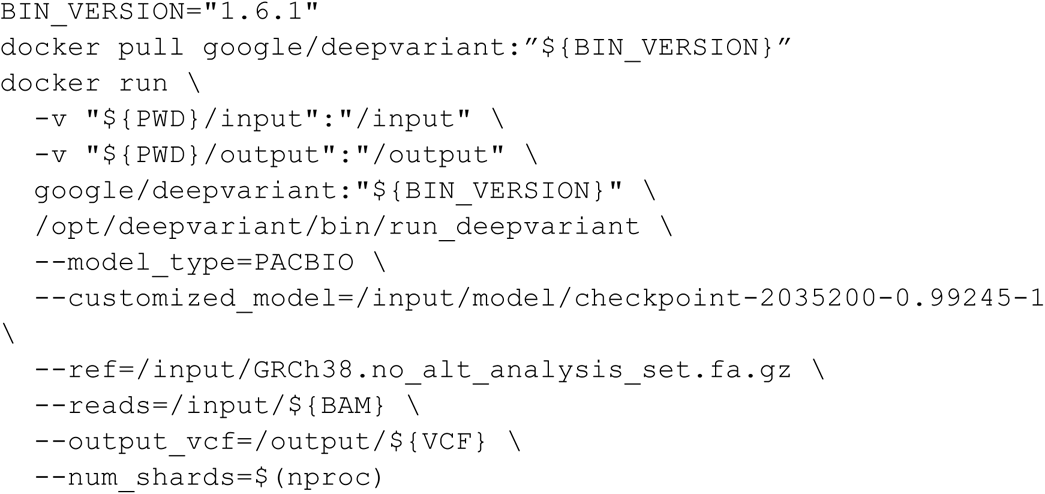

The BAM file used for this evaluation is available at (Chromosome 20 was used for all evaluations):

https://storage.googleapis.com/brain-genomics-public/research/platinum/bam/NA12878.GRCh38.haplotagged.35x.bam

https://storage.googleapis.com/brain-genomics-public/research/platinum/bam/NA12878.GRCh38.haplotagged.35x.bam.bai

The reference file is available at:

https://storage.googleapis.com/brain-genomics-public/research/platinum/reference/GRCh38.no_alt_analysis_set.fa.gz

The full number of true variant calls, errors, precision, and recall numbers on Chromosome 20 are listed for GIAB in **Table S11** and on the Platinum Truth Set in **Table S12.**

## Supplementary Figures

**Fig. S1.**
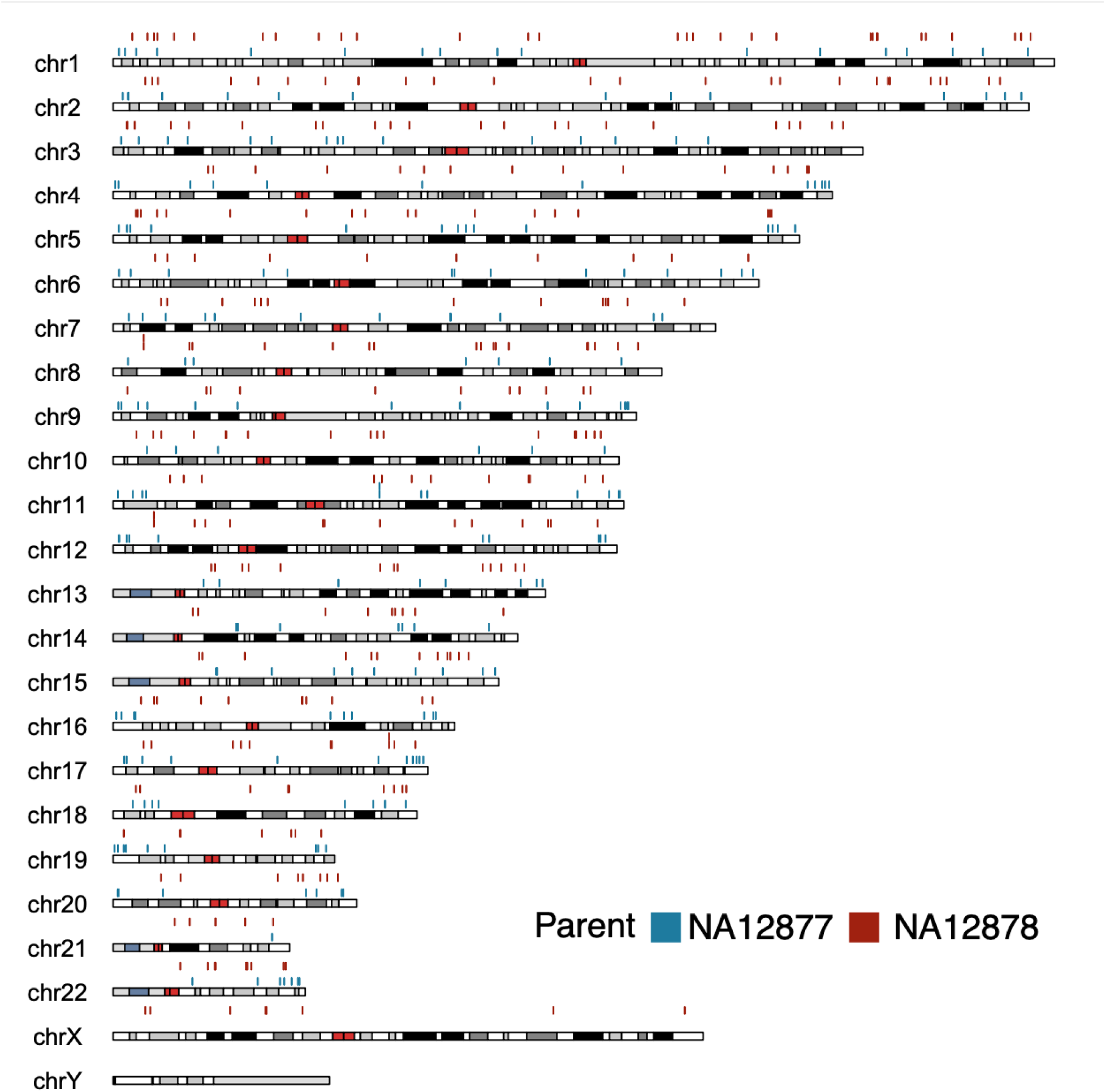
Distribution of recombination events in the third generation of CEPH-1463 as determined by inheritance vector analysis. Blue is NA12877 (paternal), red is NA12878 (maternal). Each tick mark denotes a recombination breakpoint interval.

**Fig. S2.**
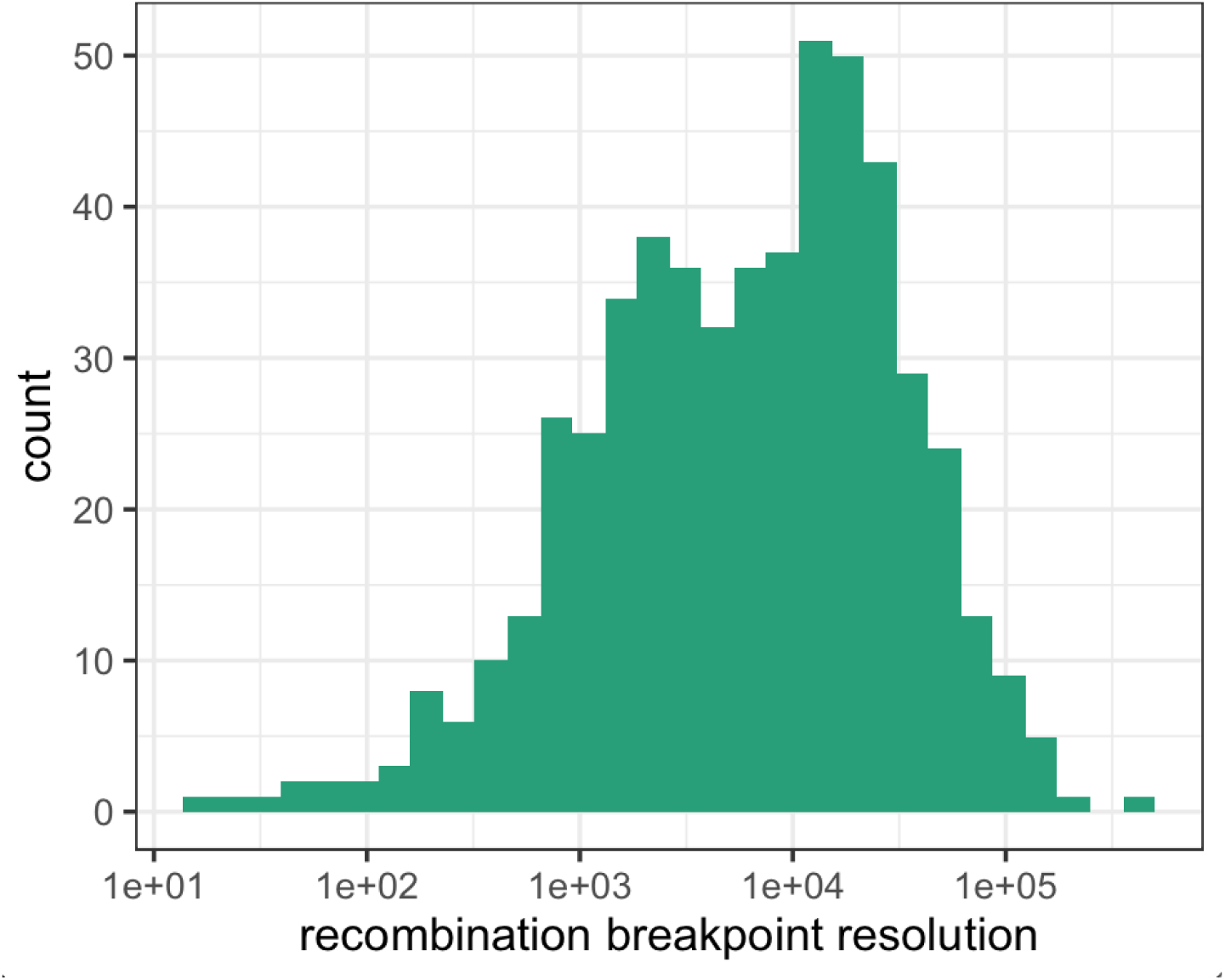
Recombination gap sizes in the inheritance vectors. The mean resolution is 17 kb and the median is 7 kb. The x-axis units are bp.

**Fig. S3:**
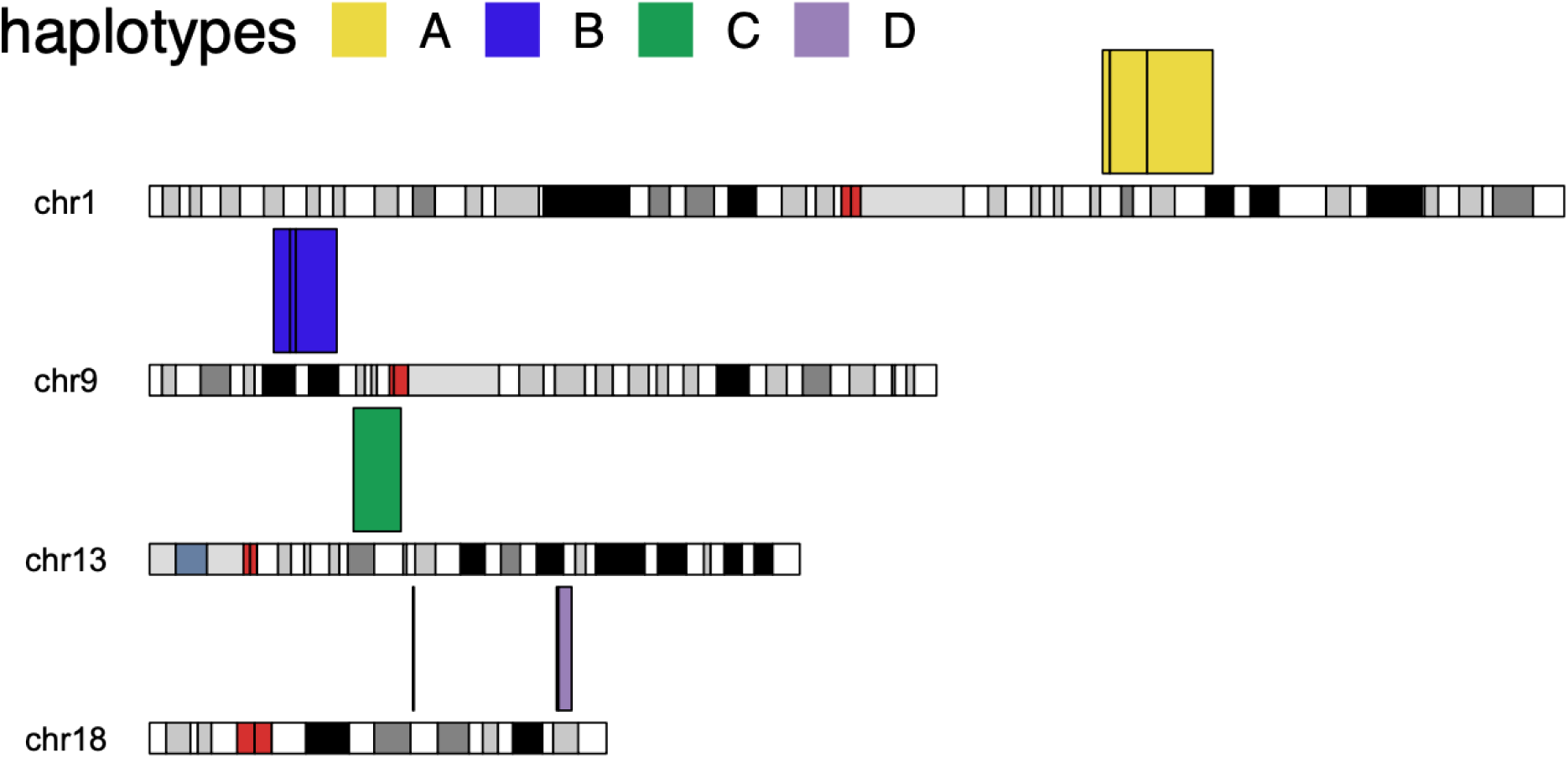
Autosomal ideogram highlighting the 41 Mb where one paternal (A/B) or maternal (C/D) haplotype was not inherited by any of the third (G3) generation family members. In total, there are ten regions.

**Fig. S4.**
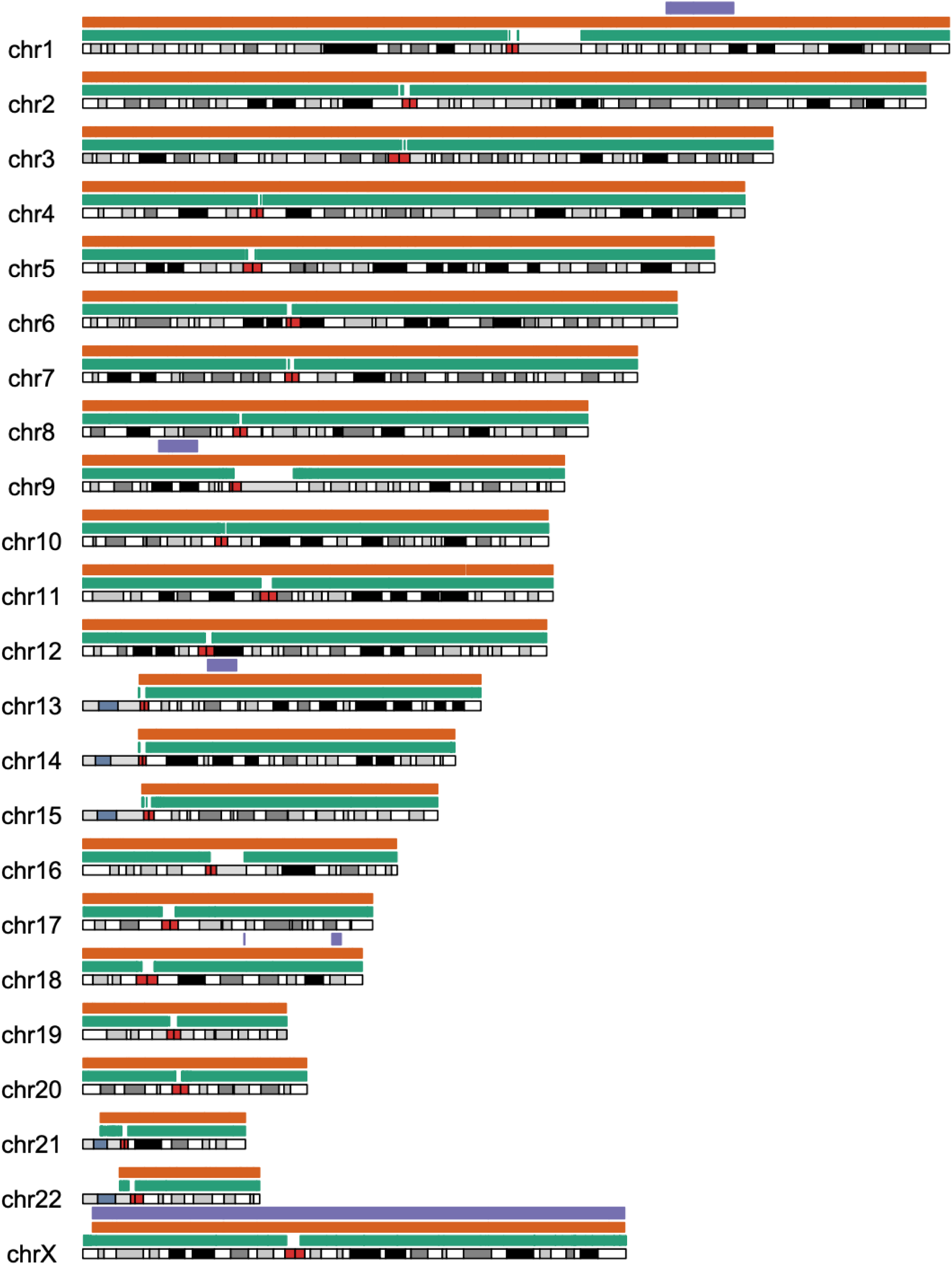
An ideogram of the coverage regions (green), inheritance vectors (orange), and missing haplotypes (purple). There are 668 regions with at least 10-fold coverage in all ten samples, there are 558 regions with defined inheritance vectors. The missing haplotype on Chromosome X is from the paternal lineage.

**Fig. S5.**
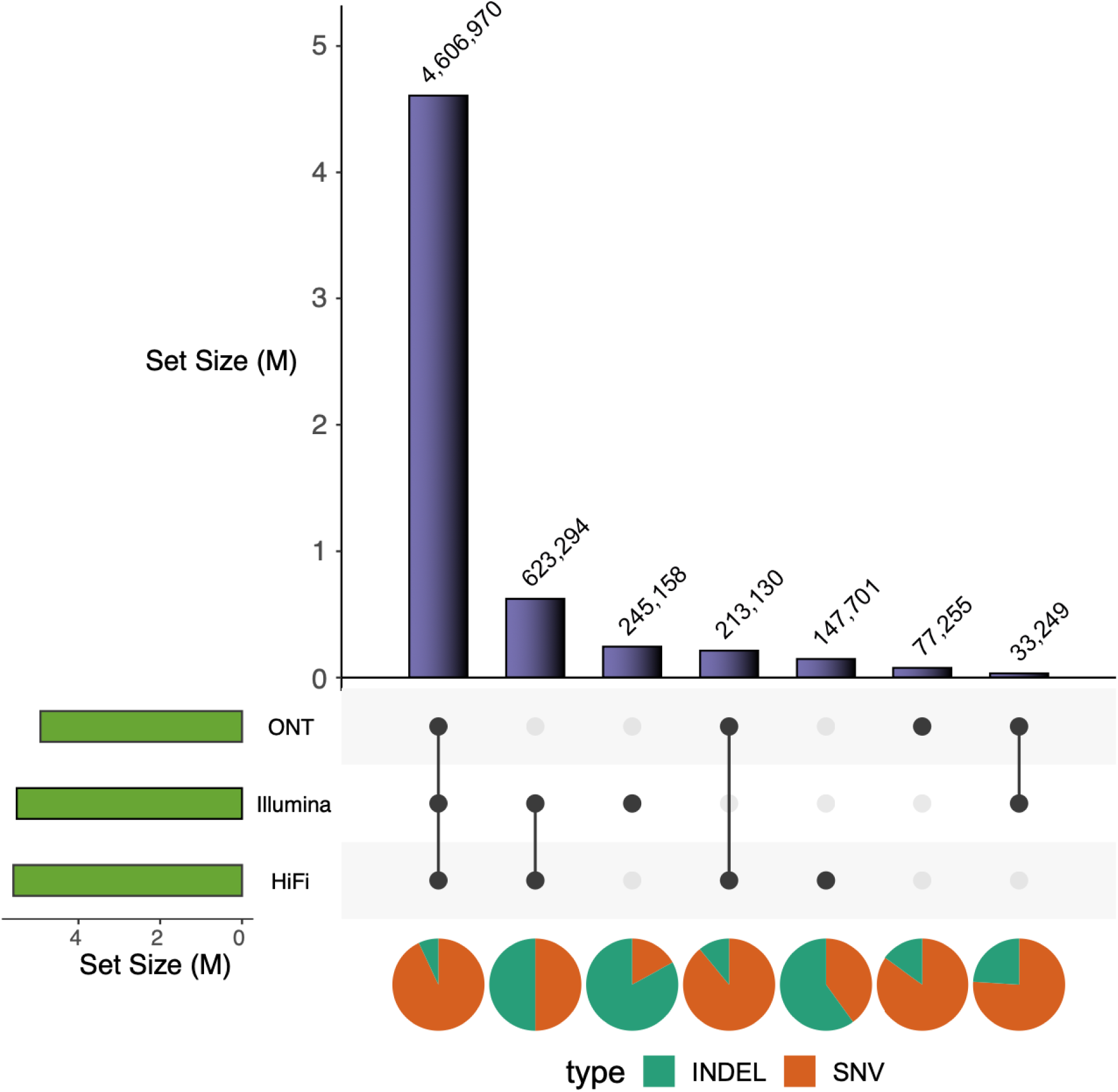
Overlap for Illumina, HiFi, and ONT data in the merged small variant truth set (SNVs and indels). The pie charts show the ratio of SNVs and indels for each category. Most variation is shared between all three sequencing technologies.

**Fig. S6.**
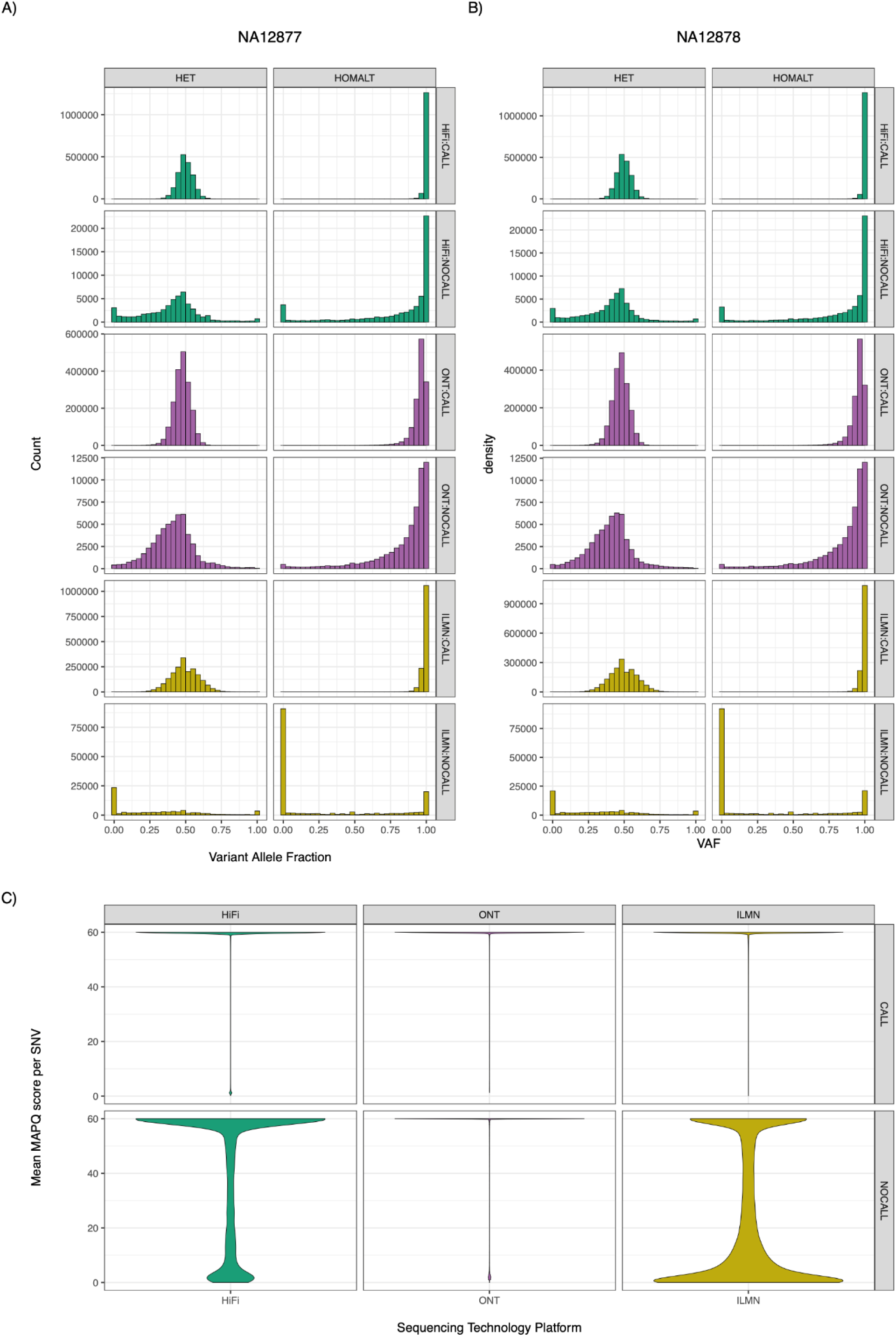
Visualizing pileup analysis. **A-B)** Histogram of variant allele frequency for approximately 4.93M pedigree-consistent SNV sites in the G2 generation. **C**) Violin plot of average MAPQ score per variant. Each figure is labeled with the sequencing platform and specifies whether the variant was pedigree-consistent and called or not called during joint genotyping. An SNV was classified as a CALL if it was identified by all three sequencing platforms. In contrast, an SNV was defined as a NOCALL for each sequencing platform if it was not called during joint genotyping and if it was not pedigree-consistent in the initial joint call. ILMN, Illumina; HiFi, high-fidelity; ONT, Oxford Nanopore Technologies; HET, heterozygous; HOMALT, homozygous alternate.

**Fig. S7.**
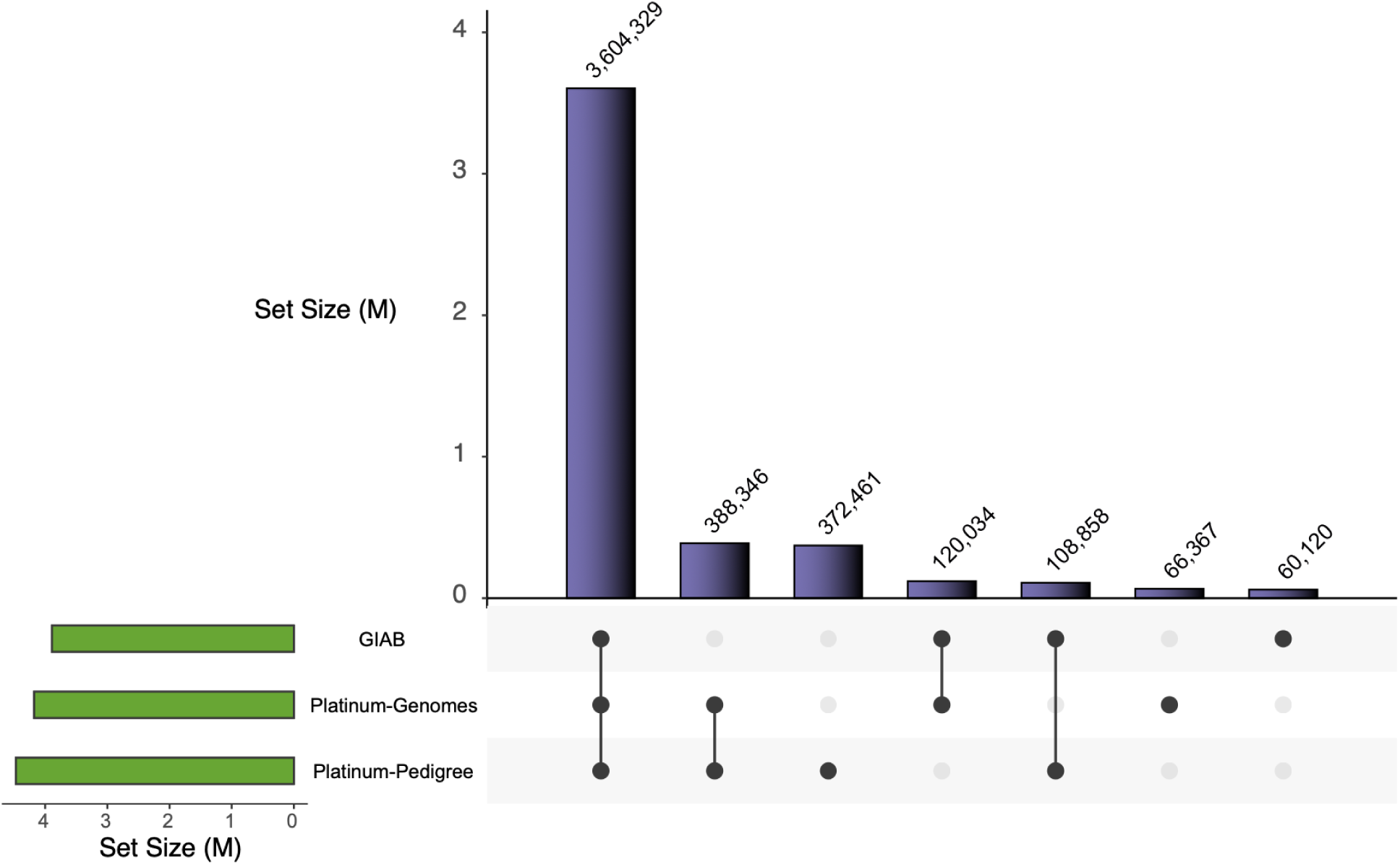
Small variant call sets overlap for GIAB 4.2.1, Platinum Genome, and Platinum Pedigree. Most variation is common to all three truth sets.

**Fig. S8.**
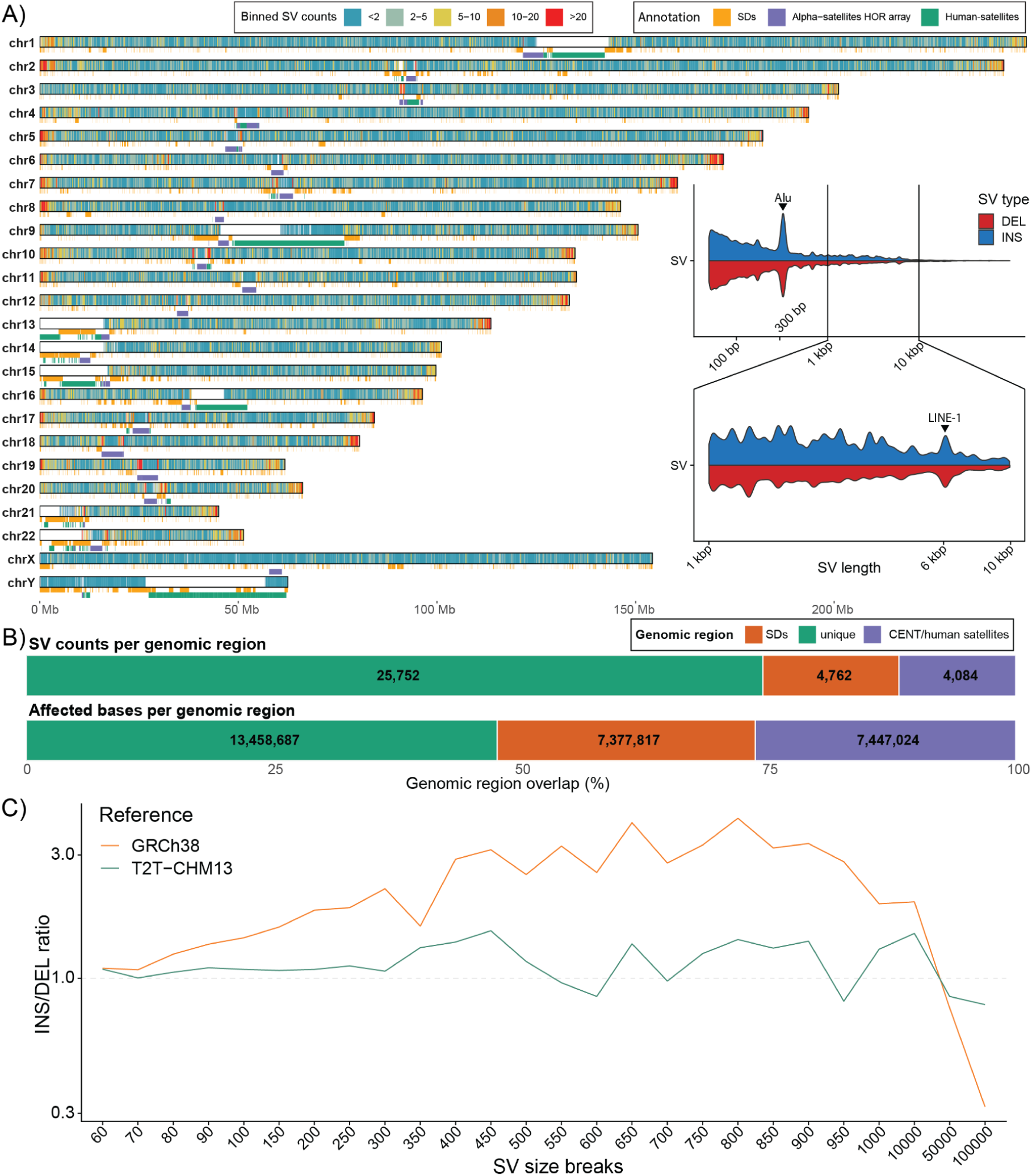
SV density across the pedigree relative to T2T-CHM13. **A)** Density of SVs with respect to T2T-CHM13. SVs are counted in 200 kb long bins. T2T-CHM13 SD annotation is shown as orange-colored rectangles below each chromosome along with the annotation of centromeric satellites shown as purple and green rectangles. Inset shows a size distribution of SVs for insertions (blue) and deletions (red) above and below the midline, respectively. We mark increased frequency of ALU elements around 300 bp as well as LINE-1 elements around 6 kbp. **B)** Top bar shows counts of SVs overlapping non-SD regions (green), SD regions (orange), and centromeric satellites (purple). Bottom bar shows the amount of affected base pairs by SVs overlapping non-SD regions (green), SD regions (orange), and centromeric satellites (purple). **C)** Ratio of insertions and deletions per defined SV size breaks (x-axis) shown separately for SV detected with respect to GRCh38 (orange) and T2T-CHM13 (green).

**Fig. S9.**
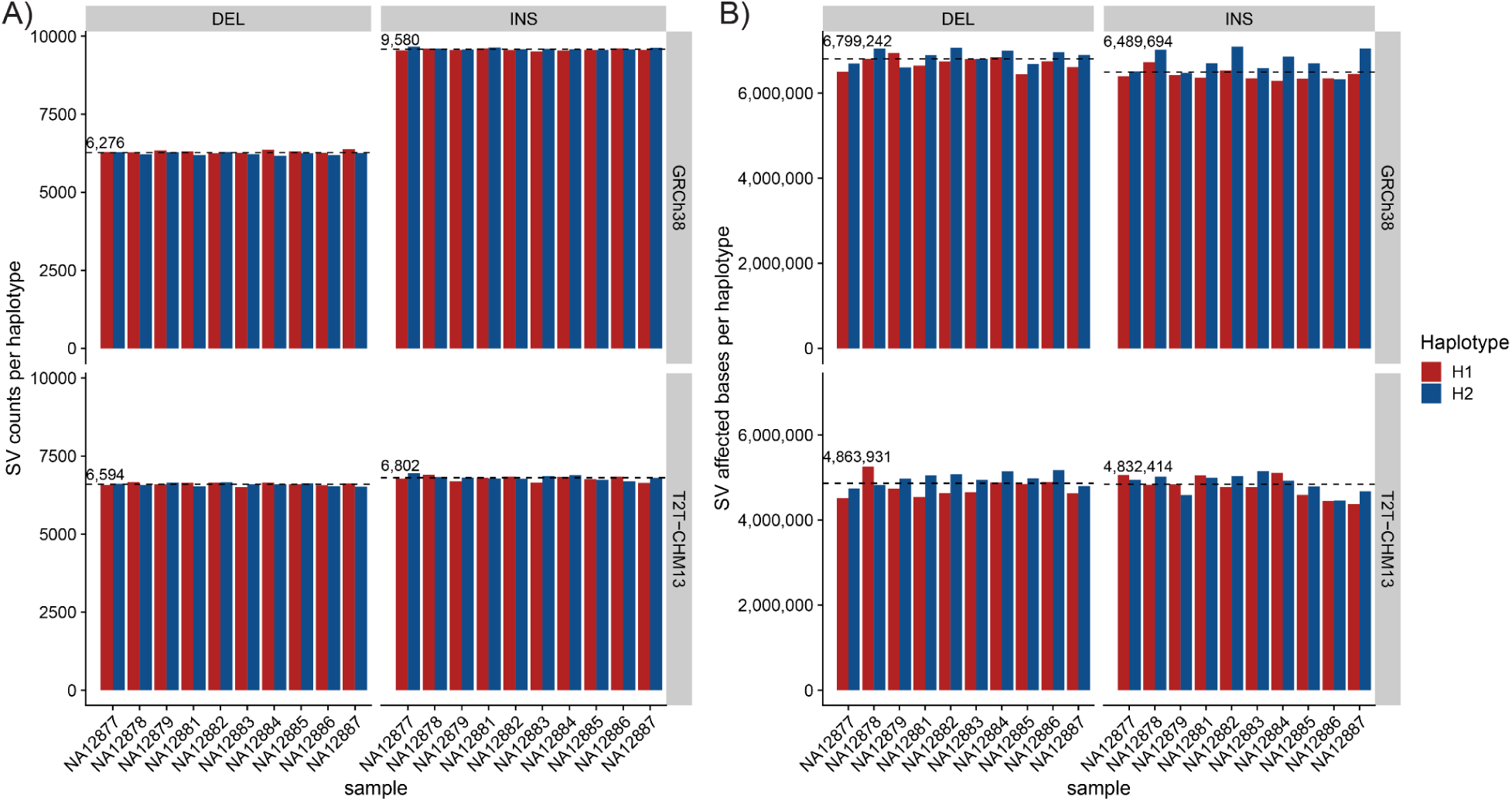
Overall number of SVs and affected bases per haplotype. **A)** Total number of SV counts per sample and per haplotype (x-axis). Counts are shown separately for insertions (INS) and deletions (DEL) with respect to both references (GRCh38 and T2T-CHM13). **B)** Total number of affected bases by SVs per sample and per haplotype (x-axis). Counts are shown separately for insertions (INS) and deletions (DEL) with respect to both references (GRCh38 and T2T-CHM13).

**Fig. S10.**
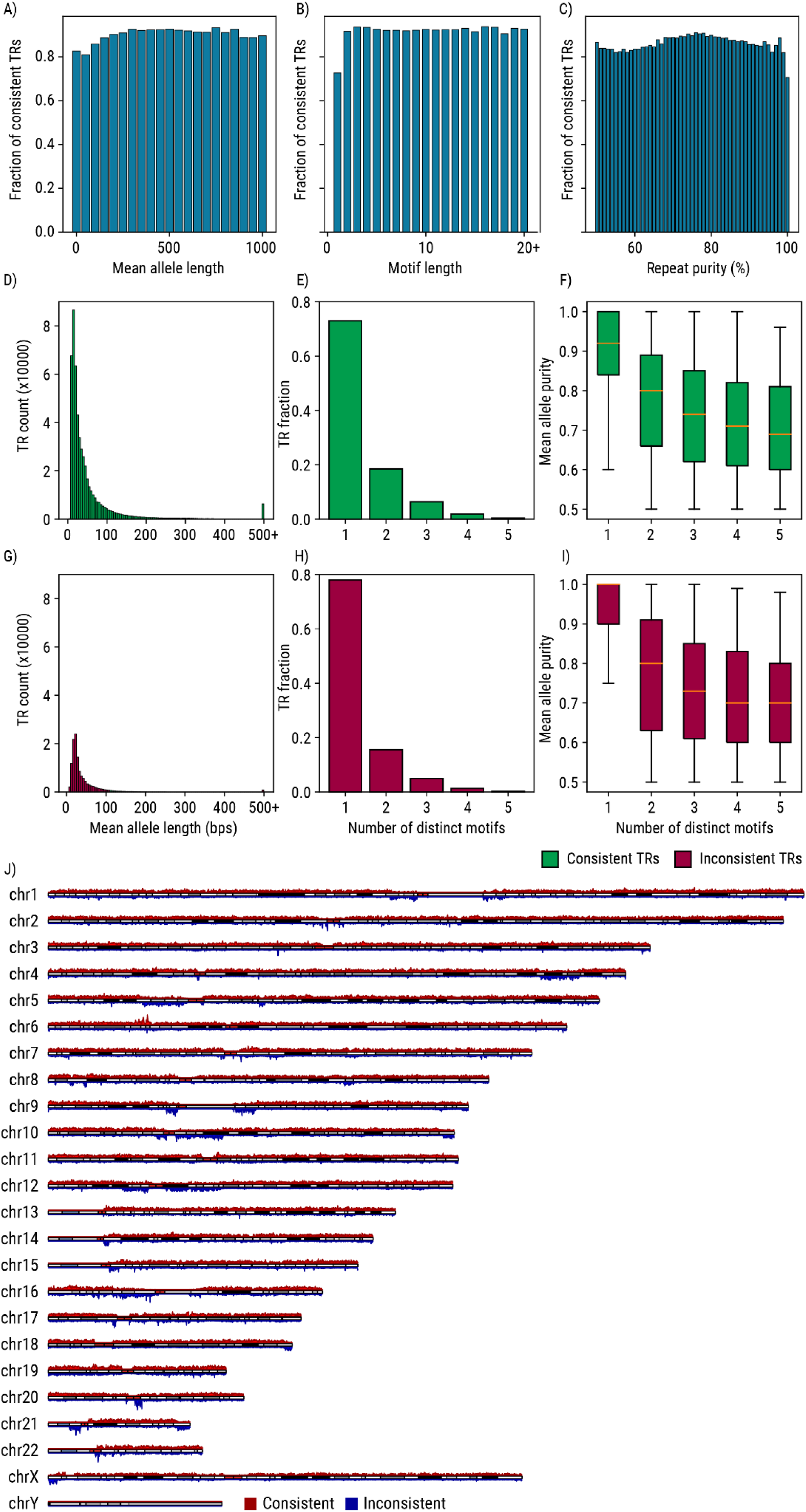
Properties of tandem repeats. Fractions of pedigree-consistent TRs stratified by the **A)** mean allele length, **B)** motif length, and **C)** motif count. **D)** Distribution of the mean allele lengths for consistent TRs. Distribution of **E)** consistent TRs and **F)** their purities stratified by the number of distinct motifs they are composed of. **G)** Distribution of the mean allele lengths for inconsistent TRs. Distribution of **H)** inconsistent TRs and **I)** their purities stratified by the number of distinct motifs they are composed of. **J)** Karyotype plot showing the distribution of genomic locations of pedigree-consistent and inconsistent TRs.

